# Ancient DNA in *Lycoptera* Fossils: Key Discoveries for Evolution and Methodological Reassessment

**DOI:** 10.1101/2023.06.18.545504

**Authors:** Wan-Qian Zhao, Zhan-Yong Guo, Zeng-Yuan Tian, Tong-Fu Su, Gang-Qiang Cao, Zi-Xin Qi, Tian-Cang Qin, Wei Zhou, Jin-Yu Yang, Ming-Jie Chen, Xin-Ge Zhang, Chun-Yan Zhou, Chuan-Jia Zhu, Meng-Fei Tang, Di Wu, Mei-Rong Song, Yu-Qi Guo, Li-You Qiu, Fei Liang, Mei-Jun Li, Jun-Hui Geng, Li-Juan Zhao, Shu-Jie Zhang

## Abstract

Obtaining high-quality ancient DNA (aDNA) is a crucial process for advancements in molecular paleontology. However, current research is restricted to a limited range of fossil types due to DNA degradation and contamination from environmental DNA (eDNA). Utilizing the technology of nanoparticle affinity beads, we successfully extracted DNA from *Lycoptera davidi* fossils and obtained 1,258,901 DNA sequences. To ascertain their origins, we developed and implemented a novel protocol, the "mega screen method", designed to categorize aDNA fragments. This approach successfully identified 243 Actinopterygii species-specific fragments (SSFs), which subsequent analysis confirmed as original *in situ* DNA (oriDNA) sequences. *Lycoptera davidi,* a ray-finned fish from the Early Cretaceous Jehol Biota, dates back approximately 120 million years (Ma). Notably, the absence of deamination in certain oriDNA and paleoenvironmental DNA (paeDNA) fragments calls into question its reliability as a definitive marker for aDNA identification, a phenomenon potentially attributable to the protective effects of fossil diagenesis. Additionally, we identified a novel mechanism, "coding region sliding replication and recombination", responsible for generating new transposase. This mechanism, distinct from conventional processes such as horizontal gene transfer or vertical inheritance from parents, can facilitate a rapid genomic diversification. The transposase CDS insertion into the GAS2 gene of *Anabarilius grahami* offers molecular insights into the genetic interactions between ancestral carp and *Lycoptera* and the swift emergence of the expansion of early fish species starting from the early Cretaceous. These findings enhance our understanding of fossil DNA preservation, lay the groundwork for an ancient species DNA database, and introduce innovative tools for exploring evolutionary relationships across geological timescales.

## Introduction

The acquisition of high-quality DNA data is fundamental to molecular paleontological research. Although scientists declare that they have successfully retrieved oriDNA from some fossils dating back approximately 1 Ma, extracting amplifiable oriDNA from most fossils continues to pose a significant challenge ^1^. A majority of the oriDNA has been lost, degraded, or crosslinked with macromolecules, making it unsuitable for use as an amplification template. Fossils embedded in lithological matrices can adsorb exogenous eDNA, with some potentially becoming embedded or subsequently replaced by newly introduced eDNA ^2,3,4^. eDNA is integrated into fossils at distinct temporal stages: during pre- or post-lithification phases (paleoenvironmental DNA, or paeDNA and in recent or contemporary periods (present-day DNA, or preDNA). Predominantly, the eDNA found in most fossils originates from bacteria, often overshadowing the signals of oriDNA ^5,6^. Additionally, studies have highlighted that oriDNA fragments frequently undergo deamination, which contributes to the degradation of the DNA strand ^7,8^.

The conditions under which fossils are buried and the mechanisms of their formation are crucial for the preservation of oriDNA. Research has shown that certain fossilized volcanic tuffs, rapidly encased by volcanic ash and formed from living organisms, still contain intact nuclei of *in situ* organisms preserved at the outlines of their remains. For instance, intact nuclear structures are commonly found in fossils such as Jurassic plants and the Jehol Biota; nucleic acid stains reveal the presence of nucleic acid material in the nucleus location ^9,10,11,12,13^. While these findings are fascinating, they also raise important questions. For instance, if the nucleus tests positive for nucleic acids, are these derived from oriDNA? Do they include long-stranded DNA sequences (>100 bp) suitable for amplification templates? Addressing these questions is key to driving the most critical technological advancements in molecular paleontology ^14^.

*Lycoptera davidi* is an extinct freshwater fish of the order Osteoglossiformes. As prominent fossils of the Jehol Biota, they hold significant biostratigraphic importance for the early Cretaceous period (J3-K1), dating back approximately 120 Ma. The fossils were preserved in sedimentary rock and formed under hydrostatic conditions when the fish bodies were quickly encapsulated by volcanic ash ^15,16^.

In this study, fossil DNA extraction was performed using nanoparticle affinity beads and sequencing of 1, 258,901 DNA fragments ^17^. The mega screen method was employed to distinguish oriDNA fragments from others, successfully identifying 243 *Lycoptera* fragments (oriDNAs). Additionally, numerous DNA fragments from ancient hominids were detected, likely originating from environmental sources and deposited within fossil voids in antiquity. Intriguingly, most of these fragments exhibit no evidence of deamination, suggesting exceptional preservation. Further analysis using amino acid sequence data revealed a self-renewal mechanism within transposase genes, termed "coding region sliding replication and recombination". This mechanism appears to underpin the rapid diversification of Cretaceous fish, offering significant insights into fish evolution. Moreover, the DNA evidence derived from non-skeletal remains introduces novel methodologies for evolutionary studies, fundamentally advancing the field of molecular paleontology.

## Results and Discussion

### I. Extracting DNA information using the mega screen method

Utilizing the mega screen method, 1,258,901 DNA fragments of diverse origins were classified; the relevant biological information was extracted. This methodology involves two steps outlined in Materials and Methods and Figure S1.

#### All sequences were divided into subsets based on the lineage

Initially, as the *Lycoptera* genome data was unavailable, we conducted a BLAST search to align the fossil DNA sequences with all sequences in the NCBI database (Version 5, Nucleotide sequence Database) using the Best E - value mode (BE mode). We processed data from the 1,258,901 reads, creating subsets based on genealogical lineages and determining the total number of sequences (TS) within each subset. Among these, 674,472 total sequences (TS) were classified as bacterial sequences, 131,312 TS as primate sequences, 30,211 TS as angiosperm sequences, and 26,852 TS as fungal sequences. Additionally, 19,747 TS were categorized in the arthropod sequence subset. With the 10,000 Fish Genomes Project (F10K) nearing completion, the genomes of representative species have all been sequenced, providing a more comprehensive reference database for fish sequence matching. Leveraging these advancements, we have identified 11,313 sequences aligning with the subset of ray-finned fish sequences (Table S1).

#### The Actinopterygii subset was dominated by aDNA

To establish the sequence selective threshold, we initially set the E-value below 1E-07 and identified the sequences that met this criterion as qualified sequences (QS). Among the screened sequences, 693 met the threshold, while 10,620 did not. We speculate that the inability of these *Lycoptera* sequences to meet the threshold could be attributed to the absence of *Lycoptera* genome data, inadequate matches, and limited similarity to contemporary fish genomes. Some unaligned sequences may originate from non-fish species that have not yet been sequenced. Due to these limitations, we can’t analyze these sequences in more detail in this work.

To assess the closeness of a subset to the respective modern genomes, we introduced the metrics, Affinity Index, and set the thresholds at 90% and 97.5%, respectively (see Materials and Methods). The percentage of the ray-finned fish subset passing the two threshold sequences was 8.51 % and 4.91%, respectively. Compare this to other subsets; for example, the two percentage values for the human subset were 94.78% and 91.56%, respectively, while the percentages for amphibians and reptiles were 15.15% and 12.12%, respectively. In addition, our analysis should introduce some other factors, such as geographical factors; the local geography is as follows: The cold climate and hilly topography of the region result in a sparse distribution of amphibians and reptiles, which limits their contribution to the local eDNA; therefore, the Affinity Index of this subset can be used as a background value to screen the corresponding lineages. From the above, it can be assumed that the ray-finned fish subset meets our criteria, and most of the sequences in this subset do not come from the local environment, which are aDNA or oriDNA. The Affinity Index for the human subset is high (Table S1), and most of the DNA fragments in this subset are recently contaminated preDNA sequences. As many extant species have yet to be sequenced, the values of the Affinity index for some subsets are correlational and may fluctuate in future studies.

#### Some features of the ray-finned fish DNA sequences

When comparing the DNA sequences of 243 ray-finned fish (Tables S3A, S4) to the sequences in the human subset, it was observed that they are similar in length and GC content. However, there were notable differences in metrics such as Per Identity, Query Cover, average number of base deletions, and QS/TS (%). The human subset displayed a lower frequency of insertions/deletions when compared to that of ray-finned fish due to the sequence differences between preDNA and aDNA. The DNA sequences within the fish subset demonstrated aDNA characteristics overall, as depicted in Table S2, providing a sound basis for effectively distinguishing the ray-finned fish subset (Subset Affinity: 52.87%; see Materials and Methods) from the preDNA-dominated human subset (Subset Affinity: 97.15%).

#### Identification of each sequence in the fish subset

In the subset of ray-finned fish, we initially utilized the Best E-value mode to identify 693 high-quality matches. However, some of these matches had the same E-value when hitting genomes of different species, which made it difficult to determine their genealogical affiliation. Therefore, we concluded that these sequences exclusively trace their genealogical origin to ray-finned fish. Consequently, we used the MS mode to ensure the reliability of the uniqueness hit, thereby enabling the identification of species-specific fragments (SSFs; Materials and Methods). Consequently, we confirmed 243 sequences as Actinopterygii SSFs.

It is reasonable to deduce that the sequences of the ray-finned fish subset can be divided into three parts: oriDNA sequences, paeDNA sequences, and preDNA sequences. By excluding paeDNA and preDNA sequences, oriDNA sequences can be recognized. These 243 sequences were then categorized into three groups: the "local fish group", consisting of 180 sequences (Table S4); and the "non-local fish group (including marine fish)," which contains 49 sequences. The third group, the "other ray-finned group," comprises 14 sequences that could not be classified at the order level, and its Affinity Index value is only 77.51%, indicating a notable distinction from the known genome.

#### The *Lycoptera* oriDNA fragments

We further determined the source of the sequences of the Actinopterygii subset by combining the influence of geographical factors, geological changes, and evolutionary laws. Since there has been no marine transgression since the Cretaceous, the 49 sequences of the non-local fish group should be oriDNA. This result also shows a genomic connection between the *Lycoptera* and modern marine fish. In the local fish group, the 180 sequences mainly belong to Cypriniform genomes. The average values of the Affinity Index of the above two groups of sequences are 80.34% and 83.85%, respectively, and their conservation levels (average values) are almost equal. In addition, the fossil site is currently a hilly area without lakes and rivers. These conditions indicate that most of the sequences in the local fish group cannot be preDNA. At the same time, considering freshwater fish eventually become prey to other species, it is difficult for their DNA to enter the environment and become eDNA. Therefore, we can roughly infer that most of the 180 sequences in the local fish group and the 14 sequences in the non-order group could be oriDNA (Tables S3, S4).

### II. Fossil DNA: The multiple resources hypothesis

The *Lycoptera* fossil can be defined as a rock that contains internal voids. We have developed a method to quantify the percentage of these internal spaces relative to the total volume of the rock, which ranges from 10.8% to 11.3% (see Materials and Methods). Due to diffusion and osmosis, various molecules not embedded in the fossil may be lost as internal liquid water drains away when the surrounding environment becomes arid. Subsequently, when water returns to the environment, DNA molecules may be transported into the dry voids within the fossil and become trapped. Furthermore, some molecules might be encased by minerals in the water and deposited onto the inner surfaces of the rock. However, if the environment dries out again, free molecules could escape.

Given that advanced primates exhibit the social behavior of burying their dead, DNA from the genomes of these primates and associated species, such as raising animals, prey, and surrounding vegetation, can continuously be introduced into the environment of long-term settlements, contributing to eDNA. Consequently, eDNA can infiltrate fossil rocks, and some of this DNA may also become buried in minerals within the water and permanently retained in the fossil. Thus, it is reasonable to conclude that eDNA from later periods can enter the internal voids of fossils formed in the Cretaceous period.

The mechanism of DNA preservation in *Lycoptera* fossils remains unclear. Water is the primary factor in DNA deamination^18^. Fossils in sedimentary rocks are formed when organisms are rapidly covered by volcanic ash in a calm water environment (Figure 2). This forms a protective shell around the organism, which prevents the chemical groups on the DNA molecules from coming into contact with water, allowing nucleic acids to be preserved for a long time and possibly protecting them from external elements such as oxygen and chemicals. Although some oriDNA fragments may be lost or degraded due to geological processes, remnants may still exist. The eDNA in fossils comes from the environment, adsorbed on rock surfaces, cavities, and cracks; part of it has become permanent sediment ^2,3,4,5^.

Our proposal delineates three sources of fossil DNA, as illustrated in Figure 2: the first being oriDNA before fossil formation, including the fish DNA and DNA from *in situ* species, such as food found in the stomachs of fishes, minute aquatic organisms, and fish parasites. The second source is paeDNA, and the third is preDNA. preDNA primarily originates from human activities, the surrounding plants, raising animals, parasites, bacteria, viruses, etc. The Beipiao area was a center of agriculture and animal husbandry in China. Ancient eDNA has infiltrated fossils since the remains were buried in mud under hydrostatic conditions, particularly during periods of considerable lacunae before the fossilization of dense formations. Following full petrification, the impermeability of shale presents challenges for water-soluble compounds to permeate the pores. Despite these, the possibility remains that minuscule quantities of matter continue to seep into the fossils.

To elucidate the origin of DNA in fossils, we developed a model diagram based on our hypotheses and speculations (Figure 1, right): (1) Before the fossil forms a dense structure, the internal DNA fragments are primarily composed of oriDNA. At the same time, paeDNA molecules gradually enter the pores and remain inside. (2) As the fossilization process progresses, the interior becomes denser, causing the voids that can accommodate foreign molecules to decrease and sometimes disappear entirely. Concurrently, the ingress and egress of internal DNA molecules are obstructed, embedding the DNA within the rock formation and rendering it immobile. (3) Following this process, the surface area of the internal pores continuously decreases, reducing their capacity to accommodate foreign molecules. The remaining open gaps are relatively robust and resistant to collapse or blockage. Subsequent generations of external paeDNA can still enter and adsorb to the surface of the pores, but are rarely embedded and remain predominantly mobile. (4) These mobile DNA fragments are displaced over time by later generations of external molecules. This replacement process persists over time, with older molecules being supplanted by newer ones up to the present day. (5) The oriDNA and paeDNA embedded within the fossils undergo continuous degradation. Consequently, this leads to a decrease in oriDNA and paeDNA and an increase in preDNA. In this study, a limited number of fish fragments (11,313 TS) were detected, whereas a larger quantity of human DNA fragments (127,977 TS) was identified, predominantly originating from preDNA (Table S1). This exemplifies the phenomenon of so-called "fewer old DNA molecules and more new ones" in fossils (Figure 1).

**Figure 1.**
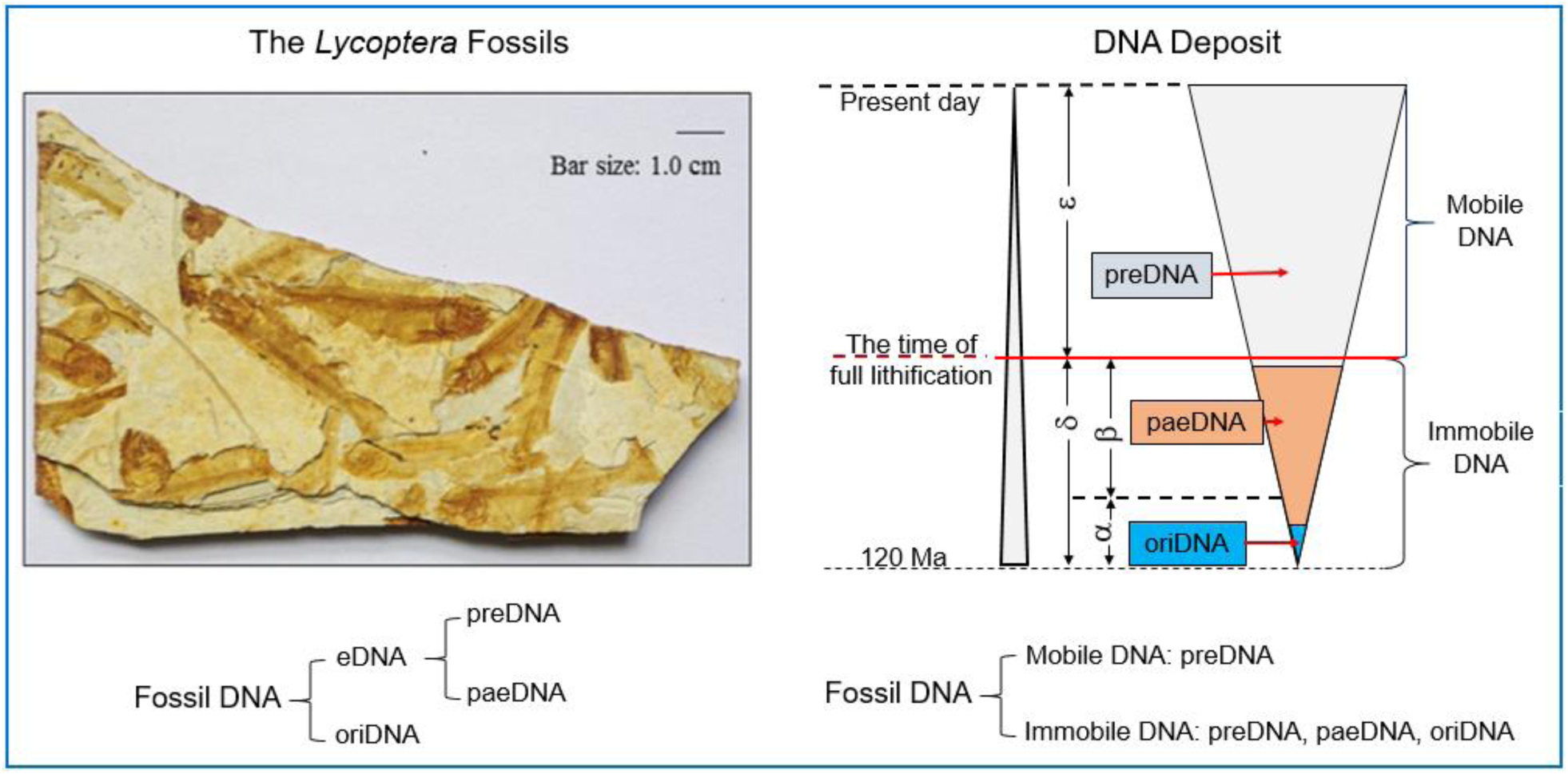
Formation of Fossils and Their DNA Contents **The fossils** were found in Beipiao City, Chaoyang City, Liaoning Province. The longitude is 120.84, and the latitude is 41.60. The process of fossil formation and preservation of DNA: During early lithification (α), a significant amount of volcanic ash descends into the water, rapidly forming a thick, viscous paste that envelops and buries the fish within a tranquil, undisturbed water environment. This results in enclosed inclusions isolated from their surroundings, allowing some paeDNA to become trapped within them. As lithification progressed (β), these inclusions gradually desiccated and were compressed into a thin, dense layer known as the “fish layer”, each approximately 1 mm thick. The brown sections in the shape of fish within a plate represent the fish remains, with the surrounding whitened sections acting as the cofferdam area. Additionally, there is the “non-residual layer” without any fish remains. After complete lithification (δ), the “fish layer” and the “non-residual layer” are mixed and overlapped in multiple layers to form dense tuff plates and are further petrified into lithological fossils. After the soft tissue has decayed, the area between the skeletal remains becomes very porous, with many tiny cavities connected to surface cracks in the fossil rock. This allows for the transfer and exchange of internal and external molecules, providing a pathway for eDNA to enter the fossil.

### III. Absence of deamination in select oriDNA and paeDNA fragments

Although deamination is a type of DNA damage and a criterion for assessing aDNA in many studies, non - deamination was observed across the 243 oriDNA sequences (Tables S3A, S4). It may be attributed to the sealed structure of fossils following diagenesis, which likely restricts exposure to external environmental conditions, thereby shielding oriDNA from outside factors. Nevertheless, some DNA fragments located on fossil surfaces, such as paeDNA or preDNA, may remain vulnerable to environmental influences, lacking the protective rock encasement and resulting in deamination. These heavily altered sequences were identified and excluded through analysis of the MS mode.

We identified numerous fragments that aligned with non-human primates’ genomes (NHP genomes) in our DNA extracts. The affinity values for these fragments range from 38% to 90%, and the mega screen method suggests a "unique" relationship between some sequences (SSFs) and the NHP genomes (Table S3B). Additionally, several facts support an opinion: these fragments likely originate from unclassified genomes of ancient Hominoidea; e.g. (1) The *Pan* lineage is not distributed throughout the East Asia region, and there are no fossil records of *Pan* species in the fossil production area, indicating these fragments are not a result of eDNA contamination; (2) our laboratory has never been exposed to samples containing *Pan* DNA. Furthermore, these fragments do not fully align with known *Pan* genomes, and there is no existing sequencing record, ruling out experimental contamination; (3) *Pan* genomes have been extensively sequenced, but the observed affinity gaps between the known ape and the sequencing data should not be so pronounced. (4) No discernible deamination was observed in these sequences.

### IV. Deamination is not suitable as the primary criterion for the authenticity of aDNA fragments

In the method developed by the Pääbo team, deamination is the main criterion for identifying aDNA fragments ^19,20,21,22,23,24^. However, several points challenge this perspective: (1) The assumption that the fossil oriDNA originates from a single individual of the corresponding species overlooks the potential for eDNA contamination from either the same or different species over time following the formation of the fossils; (2) The selection process involves only a limited number of closely related genomes as BLAST references, failing to consider the broader influence of numerous environmental species and neglecting to utilize all available genome sequencing databases; (3) There is insufficient consideration of the unreliability of BLAST results, mainly when the E-value is excessively high, rendering them outside a credible interval; (4) In this work, many DNA fragments with non-deamination match to the genomes of ancient organisms, including *Pan troglodytes*, suggesting that this characteristic does not serve as a reliable logical criterion for oriDNA identification (Table S3A, S3B). Consequently, the genomes of Neanderthals, Denisovans, and several species assembled by Pääbo’s team likely represent "mixed genomes". The belief that deamination is necessary for identifying aDNA/oriDNA is incorrect, which has hindered the search for long-chain aDNA or oriDNA, which could offer more comprehensive genetic information. As a result, several significant flaws in the literature concerning Pääbo’s research on ancient species should be addressed, and these works should be re-evaluated.

### V. Genomic links between the *Lycoptera* and other fishes

The relationship between the genomes of ray-finned fish and their morphological development is not fully understood. The classification of these fish is mainly based on their physical characteristics. A study by Zhang concluded that *Lycoptera* belongs to the order Osteoglossiformes ^25^. Early researchers, such as Cockerell and Berg, suggested that species of the Cypriniformes order originated from Jurassic Lycoptera ^26,27^, while Rosen proposed that they may have originated from Gonorynchiformes^28^. The oldest known carp fossils date back about 60 million years. Carp fishes are found in freshwater environments worldwide, except Antarctica, South America, and Australia, indicating that their origins predate the breakup of the Pangaea continent. Molecular paleontological studies have provided valuable insights into the origins of carp. Tao’s research suggests that the genomes of carp fishes can be traced back to the early Jurassic period, around 193 Ma ^29^. Over 75% of the more than 180 sequences of *Lycoptera* oriDNA in the present study align with the carp genome, indicating a genomic connection between them. Furthermore, the genetic relationship of *Lycoptera* with carp and other native fishes requires further investigation. Although the total length of the oriDNA obtained in this study is limited, further exploration is necessary for a comprehensive phylogenetic analysis.

### VI. Decoding fossil DNA: Revealing parasites and prey

In the fossil DNA, the ID: 282 sequence (Table S3) corresponds to the 28S rDNA sequence of the class *Ichthyosporea*. These are flagellated microorganisms that belong to the Opisthokonta (unranked). This small fish parasite’s taxonomic position lies between animals and fungi ^30^. Additionally, we have identified 4 DNA sequences corresponding to the *Macrobrachium nipponense* genome (IDs: 285-287) with a Subset Affinity value of 83.24%, of which 3 DNA sequences were highly divergent from the genomes of existing species. Two other sequences corresponding to *Penaeus vannamei* and *Rhynchothorax monnioti* (sea spider) genomes, both non-native species, were hypothesized to be derived from close relatives in their ancient decapod relatives (Table S3A, ID: 283, ID: 289). Upon using 3D X-ray computed tomography to scan the fossils, we found no remains of other animals. This result suggests that these fragments originated from the parasites and prey of the *Lycoptera* and that they are oriDNA rather than eDNA (Figure S1). It also brings new insights for studying their evolution.

### VII. New mechanism for generating transposase-encoding sequences

Transposons constitute over 40% of the fish genome, with transposase as a critical component in the transposition process ^31,32^. According to Frith’s research, transposon sequences in modern genomes can be regarded as "molecular fossils," exhibiting substantial sequence similarity across diverse organisms, including prokaryotes, arthropods, and mammals ^33^. Transposons play a key role in introducing new DNA sequences into the genome. Our analysis of 10 fish aDNA fragments has revealed a novel mechanism for transposase gene formation, designated "coding region sliding replication and recombination". This mechanism diverges from established conventional processes such as horizontal gene transfer, vertical inheritance, and gene or chromosomal duplication ^34^.

This mechanism comprises three distinct types of coding: (1) The initial sliding duplication takes place within a CDS to generate a segment, followed by a shift in the reading frame that produces a different segment. Subsequently, the two distinct segments can be integrated with the segments derived from other transposase genes to construct a novel CDS (Figure 2A). (2) After sliding replication, different segments are generated, which are then assembled in a tandem manner and then connected to the CDS segments from the other transposase to form a new enzyme ( Figures 2B, 2C). (3) Indel events occurring at a single locus: when an insertion or deletion of a base in a CDS results in a shift in the reading frame. Figure 2D suggests a “deletion-correction mechanism” that can utilize the mutant CDS where the Indel has occurred to generate a novel enzyme.

**Figure 2.**
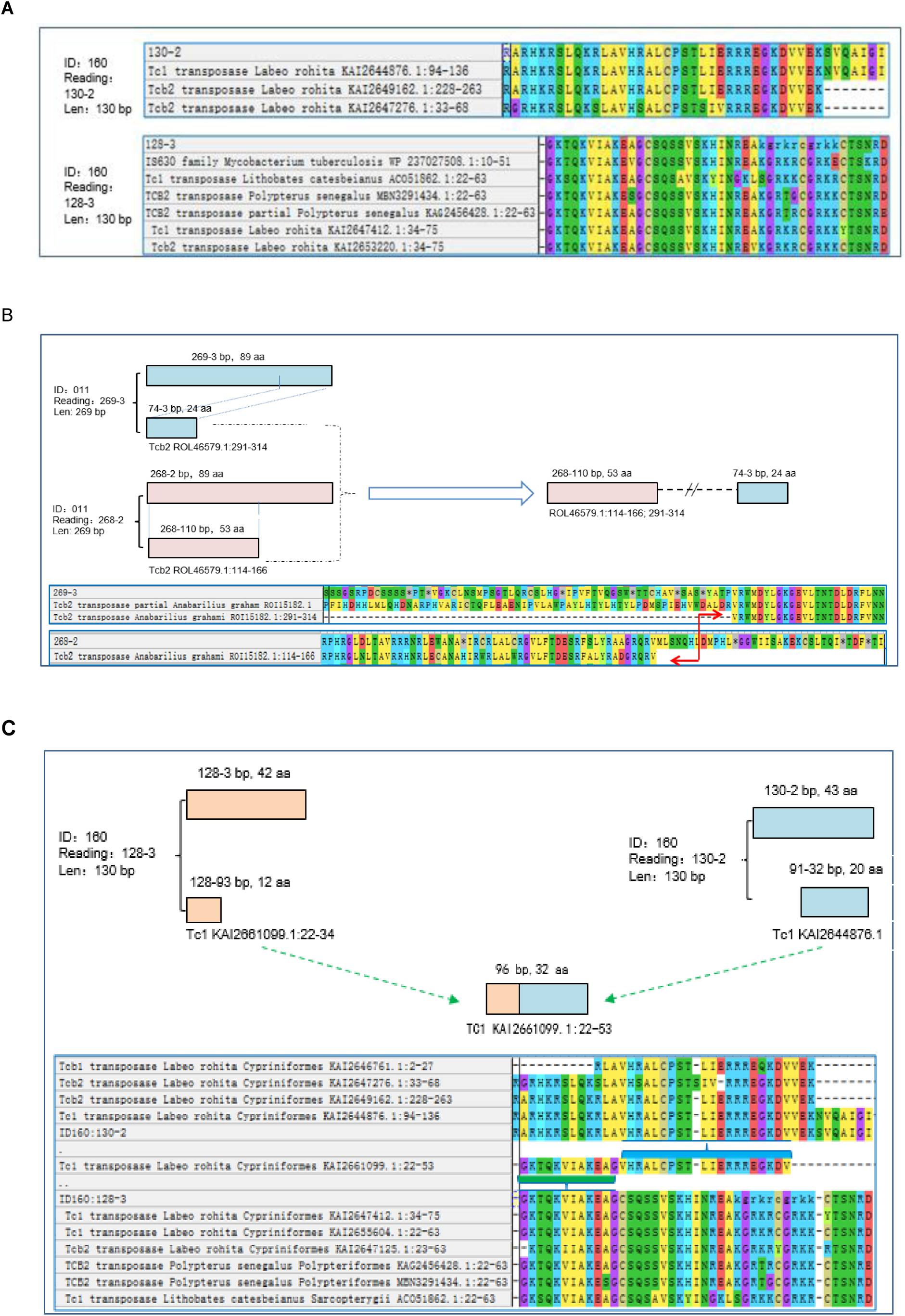
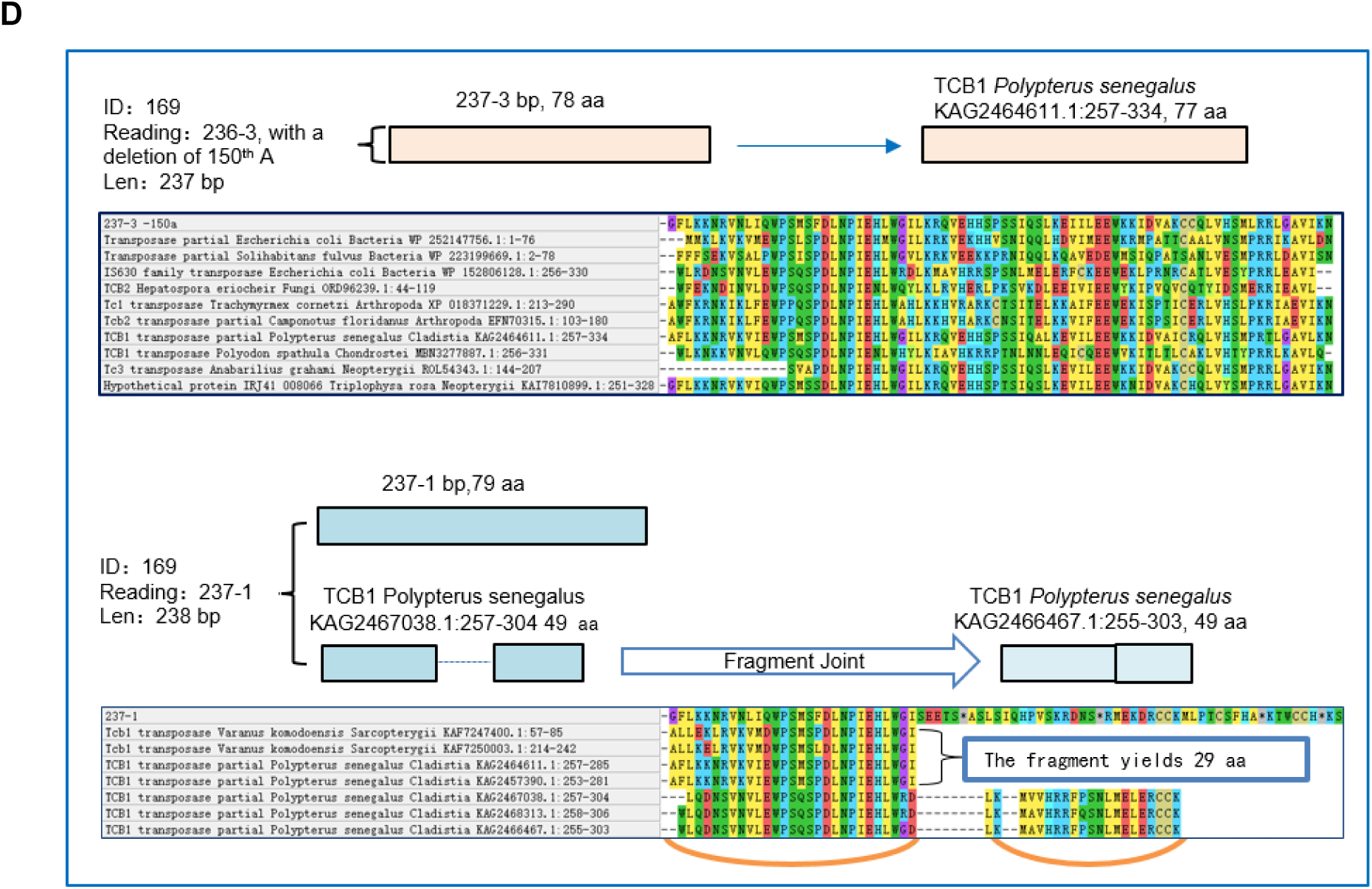
Proposed model for a new mechanism of generating transposase-encoding sequences **A**. Multiple fragments produced by sliding replication are spliced and fused with other gene fragments separately to form a new enzyme. **B**. The two fragments generated by sliding duplication were spliced together with other gene fragments to constitute the coding region of a new enzyme. **C**, the two fragments generated by sliding duplication are joined together and spliced with other gene fragments to form the coding region of a new enzyme. **D**, A site in the coding region of the transposase undergoes an Indel, but still passes clips to constitute the coding region of the new enzyme.

During the Mesozoic period, ray-finned fish underwent significant evolutionary developments, resulting in the emergence of numerous new Orders. This diversification was facilitated by the ability of their genomes to generate a substantial number of novel transposases, through the mechanism of "coding region sliding replication and recombination", independent of external DNA contributions. This intrinsic capability likely drove a rapid expansion and divergence of their genomes, accelerating the appearance of new species. Furthermore, transposases were found to integrate into the coding regions of functionally regulated proteins. For instance, the N-terminal portion of an oriDNA fragment (sequence ID: 011) shares 52 amino acids with both the Tcb2 protein and the growth arrest-specific protein 2 (GAS2) of *Anabarilius grahami*, as its C-terminal portion corresponds to the conserved region of GAS2 (Figure 3). These findings indicate that the coding DNA sequence (CDS) of the transposase was inserted into the GAS2 gene, with this translocation likely occurring before fossilization in the Early Cretaceous. This event underscores the early evolution of fish genomes and elucidates a pathway for genome content expansion. Furthermore, it provides molecular evidence supporting genetic exchange between ancestral carp and *Lycoptera*.

**Figure 3.**
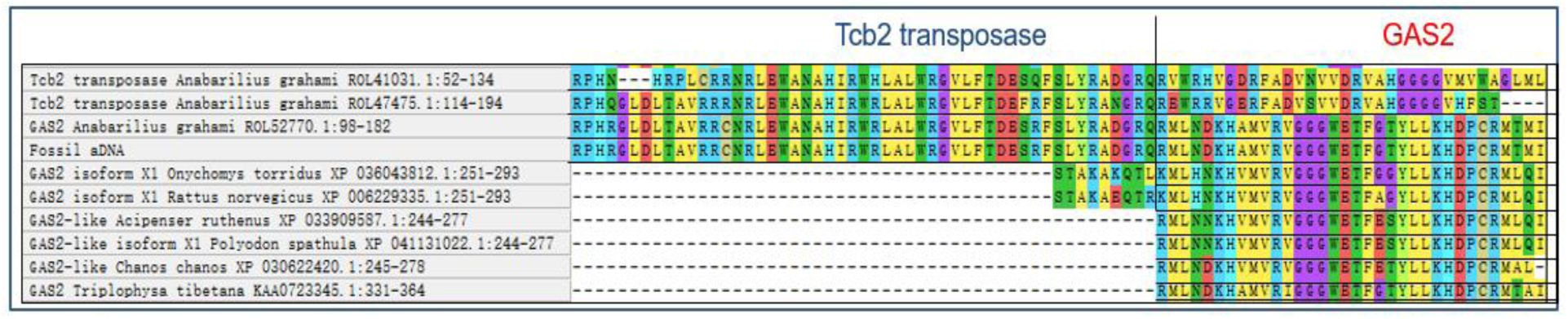
The transposase sequence is fused in the GAS2 coding region

### VIII. Summary

We developed a novel protocol, the mega screen method, employing SSF (rather than assembled genome approaches) to identify and extract information from aDNA fragments of extinct species preserved in fossils. Using this approach, we successfully identified 243 oriDNA fragments from *Lycoptera davidi*, dating to the Early Cretaceous period, approximately 120 Ma. By leveraging aDNA sequence characteristics, we traced the phylogenetic descendants of the *Lycoptera* genome, which elucidated the molecular mechanisms underlying its rapid evolution, including a novel mechanism of transposase gene formation. This intrinsic dynamic process is closely linked to the rapid emergence of new ray-finned fish species during the Cretaceous period. These findings deepen our understanding of fossil DNA composition, lay the foundation for establishing a DNA database of ancient species, and provide valuable reference points for investigating evolutionary relationships between ancient and modern species. Moreover, they enable phylogenetic tracing and exploration of molecular mechanisms driving evolution or extinction in all extinct species, significantly enriching the paleontological information landscape and opening potential pathways for the resurrection of extinct organisms.

Surprisingly, this research provides a new perspective on "Panspermia," the theory for exploring the origin of Earth’s life. Inspired by the study of extraterrestrial nucleobases in the Murchison meteorite ^36^, we propose the possibility that if rock fossils, such as the *Lycoptera* fossils, are transported into space, the DNA inside could potentially remain dormant, effectively and uniquely preserving life. Until these fossils land in a suitable extraterrestrial environment and release the contents to become the seeds of life, catalyzing the colonization of the planet.

## Acknowledgments

We extend our gratitude to the "Chao-Xiang Talent Program" foundation of Haining City, the support that covers the costs of fossil collection, DNA extraction, and DNA sequencing from 2020 to 2021. We extend our heartfelt appreciation to the following individuals for their invaluable contributions to this project: Dr. H. J. Zheng from the Shanghai Institute for Biomedical and Pharmaceutical Technologies, Dr. J. Y. Ma from the Nanjing Institute of Geology and Paleontology, at the Chinese Academy of Sciences, Dr. A. M. Bailleul from the Institute of Vertebrate Paleontology and Paleoanthropology, at the Chinese Academy of Sciences, and Dr S. P. He from the Institute of Hydrobiology, also at Chinese Academy of Sciences. Their stimulating discussions regarding the manuscript have greatly enriched our work.

## Author contributions

The study was conceptualized and designed by W.Q.Z. and Y.Q.G. The fossils were collected and overseen by F.L. and W.Q.Z. The DNA extraction from the fossil was carried out by W.Q.Z., C.Y.Z., and J.Y.Y. Alignment between the sequences and present genomes was completed by Z.Y.G. and M.J.C. DNA laboratory analysis, interpretations, taxonomic profiling, and annotation were conducted by Z.Y.G., Z.Y.T., L.J.Z., S.J.Z. and W.Q.Z. Statistical analyses were performed and completed by G.Q.C J.H.G, and X.G.Z. Phylogenetic analyses of mitogenomic DNA sequences were performed by Z.X.Q., W.Z., C.J.Z., M.F.T., and D. W. under the supervision of G.Q.C. and W.Q.Z. Figures were designed and finished by F.L., L.Y.Q., T.F.S., and M.R.S. Project coordination was managed by T.C.Q. W.Q.Z., Z.Y.T., M.J.L., and L.Y.Q wrote the manuscript.

## Data and materials availability

All data is available in the main text or the supplementary materials.

## Supplementary Materials

### Materials and Methods

#### I. The experimental procedure for the wet lab

##### 1. X-ray 3D photography

The fossils originated from Beipiao City, Chaoyang City, Liaoning Province. They underwent a cleaning process involving tap water, deionized water, and precise cutting. Subsequently, the complete fish fossil underwent 3D X-ray scanning using a Zeiss Xradia 620 Versa scanner (Carl Zeiss AG, Germany) to ensure the absence of co-deposited species. The initial scan was conducted at a resolution of 45 μm to capture the overall structure. Specific areas such as the head, thorax, tail, and periphery were scanned at a higher resolution of 6 μm for detailed examination. The ZEISS 3D Viewer software was used to examine and reconstruct 2D orthogonal slices of the scans, helping to identify any additional species present in the scans and reconstructions.

##### 2. Method for evaluating the internal volume ratio of fossils

After cleaning the fossils, dry them in a dryer (56°C, 72 hours) and weigh them (W1). Immerse the fossils in pure water for ultrasonic treatment (10 W, 2 hours). After taking them out, quickly wipe off the attached water and weigh them (W_2_). Immediately put the wet fossils in a graduated container and immerse them in water. Calculate the total volume of the fossils (V_2_) by reading the increased scale.

Internal volume of fossils (V_1_) = (W_2_ - W_1_) × density of water
Internal volume ratio of fossils = V_1_/ V_2_ × 100%

The internal volume ratios of four *Lycoptera* fossil samples are 10.8%, 10.4%, 11.3%, and 10.8%.

##### 3. DNA Extraction, DNA Library Construction, and Sequencing

The procedure was conducted in a BSL-2 industrial laboratory (Biosafety Level 2), following rigorous ancient DNA extraction protocols ^37^. The process involved using UV light to disinfect all used items. We also applied Geobio® DNA Remover (from Jiaxing Jiesai Biotechnology Co., Ltd., Jiaxing, China) on clean benches and solid equipment. It is important to note that this laboratory has never been exposed to fish, non-human primates, or plant samples before this work.

The stone was cleaned with tap water and deionized water, and the surface was clean. Immerse it in Geobio ^®^ DNA Remover for a few seconds and then quickly remove it and soak it 3 times in deionized water. The stone was washed with deionized water to remove the additional detergent solution. After drying, the stone was cut into 10 × 10 × 5.0 (cm ^3^) raw materials, and the multilayered stone was divided into single layers approximately 0.8-1.2 mm thick. The textured portion of the fish was scraped off, and the dropped particles were collected in a clean tube. Scrape off the contoured portion at least 1.0 cm from the textured portion and collect the dropped particles in a separate tube. Harvest the same weight of fine powder from the non-textured (coffered) portion of the fossil at least 1.0 cm from the textured portion. Use a set of mortar and pestle and a mortar and pestle to grind the small particles into a fine powder. Fossil DNA was extracted from the above fine powdered material using the Geobio^®^ DNA (Solid State Matter) Extraction Kit (Jiaxing Jiesai Biotechnology Co., Ltd., Jiaxing, China) ^17^.

During long-term geological evolution, some *in situ* DNA molecules within the fossils undergo cross-linking reactions with other macromolecules (e.g., proteins and polysaccharides) as a result of the action of physicochemical and biological factors, to assess the concentration of the extracts, we examined the optical density (OD) values of the DNA mix, which contains both DNA-linking complexes and free DNA molecules, at the peak of the UV 260 nm. Five replicate extractions were performed on the textured portion of the sample, and two replicate extractions were performed on the non-textured portion, the so-called cofferdam portion (Table S3).

The DNA extracted was repaired using NEB PreCR ^®^ Mix (M0309S). Next-generation sequencing (NGS) DNA libraries were prepared following the SSDLSP standard program (Sangon Biotech, CORP). Sequencing was conducted on an Illumina NovaSeq 6000 (S-4 kit) using the "Dual Label Sequencing" standard procedure. The reads met quality control standards and exhibited a satisfactory quality distribution. Occasionally, it may be necessary to manually remove excess adapters from both ends of the reads (sequences) to obtain accurate information. Combining forward reads with their corresponding reverse reads resulted in 1,259,658 paired sequences for the fossil-textured portion (including 757 unnamed sequences) and 635 sequences for the fossil non-textured portions (matrix).

#### II. The experimental procedure for the dry lab

##### 1. The mega screen method comprises two modes

Initially, we utilized the BE mode to cluster sequences by aligning fossil DNA with known sequences from the NCBI database (Version 5, Nucleotide Sequence Database). An E-value threshold of less than 1E-07 was applied to filter for qualified sequences (QS) and determine the best match at a specific taxonomic level.

Subsequently, we employed the MS mode to perform a search that excluded the initial result. If the quotient of the E-value of the first result divided by the subsequent one is ≤ 0.01 (1e-02), the fragment is considered specific to the first result (in a taxonomic unit), thereby confirming its "specificity". This indicates that the test sequence (query) corresponds to the subject identified in the initial search, suggesting a unique origin, such as Actinopterygii SSFs. Conversely, if the quotient surpasses this threshold, the sequence is presumed to be shared between two taxa, suggesting it does not derive from a single lineage. Consequently, we discarded the test sequence without deriving further conclusions.

In addition, the mega screen method can be applied to uncover relationships among similar sequences found across various genomic organisms, offering valuable insights into patterns of genome evolution.

**The whole process involves the following steps**

1. Nucleotide Blast is employed to align sequences to the entire NCBI database without any limitation.
2. Creation of subsets for sorting the sequences: Using the BE mode, results are matched, and sequences are grouped into the appropriate subsets, determining the total number of sequences (TS). Subsequently, an E-value threshold of < 1E-07 is established to filter for qualified sequences (QS).
3. The subset primarily containing ancient DNA (aDNA) is selected.
4. Each sequence within this subset is screened individually using the MS mode to identify those originating from a single lineage, categorizing the “unique” sequences (SSFs).
5. Considering various factors, including local climate shifting, geological changes, and evolutionary principles of the host species, we can ascertain if the sequence is oriDNA.

##### 2. Sequence affinity (Affinity)

This metric measures the similarity between subject sequences and hit genomes. It is calculated by multiplying the values of Identity and Cover obtained from NCBI Blast and then converting the result to a percentage (Identity × Cover × 100%). Mutations in the genomic and mitogenomic sequences occur continuously during the evolutionary process, leading to significant sequence differences between ancient and modern species. However, this variation is less pronounced in conserved sequences compared to non-conserved sequences. A high-affinity value indicates that the sequence is highly coherent and related to the genome of the modern species, which is likely to be either a conserved sequence or a modern eDNA. A low-affinity value indicates a low similarity, and the sequence maybe from a distantly related species either has or has not yet been sequenced. It is necessary to consider various factors, including the sequence composition and conservation, host species information, host genome sequencing data, etc., rather than relying on the Affinity value alone when determining whether a sequence is an oriDNA.

##### 3. Subset affinity

This metric refers to the average of the sequence affinities of all sequences in a subset; it indicates whether the subset consists primarily of aDNA sequences. High values (close to 100%) indicate that the DNA sequences in the subset consist mainly of preDNA.

##### 4. The percentage threshold for subset affinity (Affinity Index)

The number of sequences within a subset that exceed a certain threshold as a percentage of the total number of sequences. In this study, we set the thresholds at 90% (indicating limited similarity between the sequences and their hit genomes) and 97.5% (indicating high similarity between the sequences and their hit genomes), respectively. This metric reflects the closeness of the relationship between all sequences within a lineage subset and the modern genome. For a subset, a low value indicates that the sequences are mostly ancient (aDNA), while a high value indicates that the sequences are mostly recent (preDNA).

---

**Figure S1.**
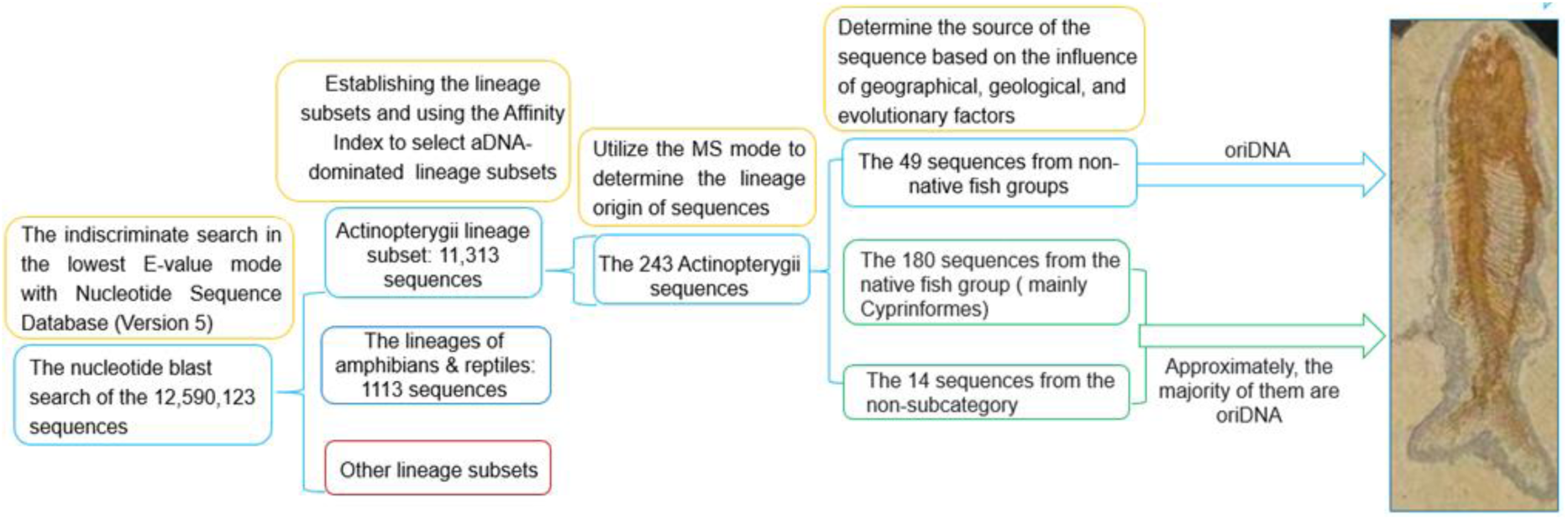
Selecting for the fossil oriDNA using the mega screen method

**Figure S2.**
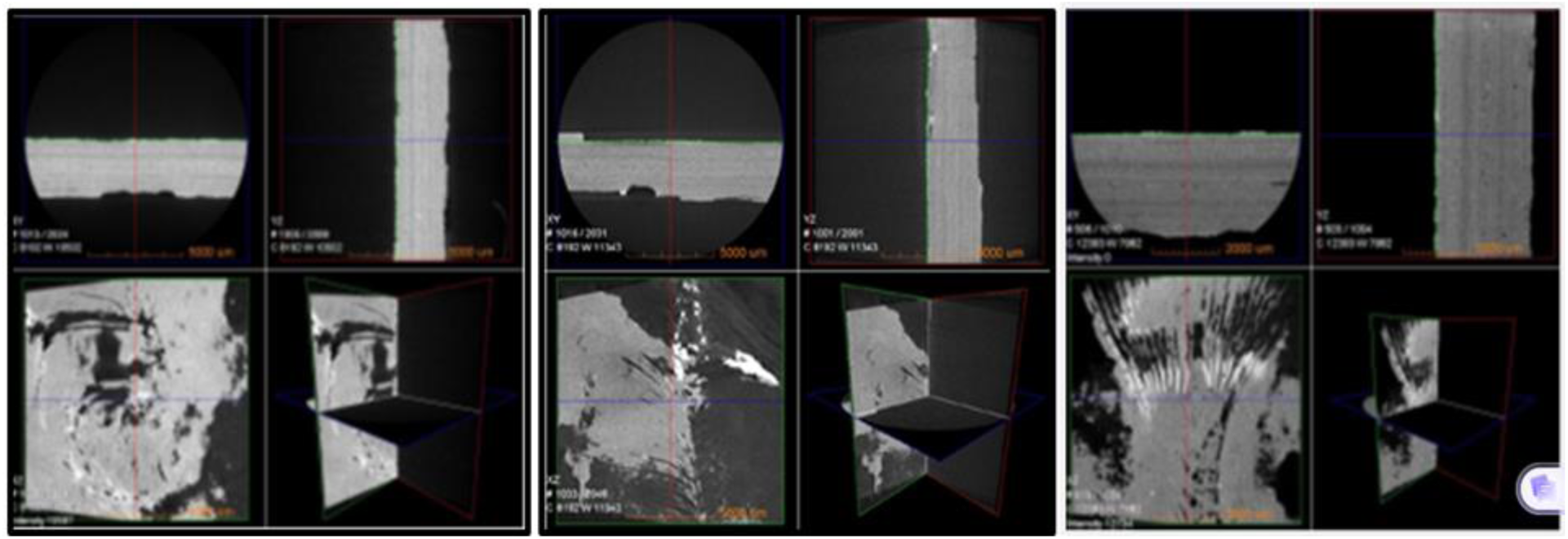
3D X-ray scanning the Fossil Fish

**Table S1.**
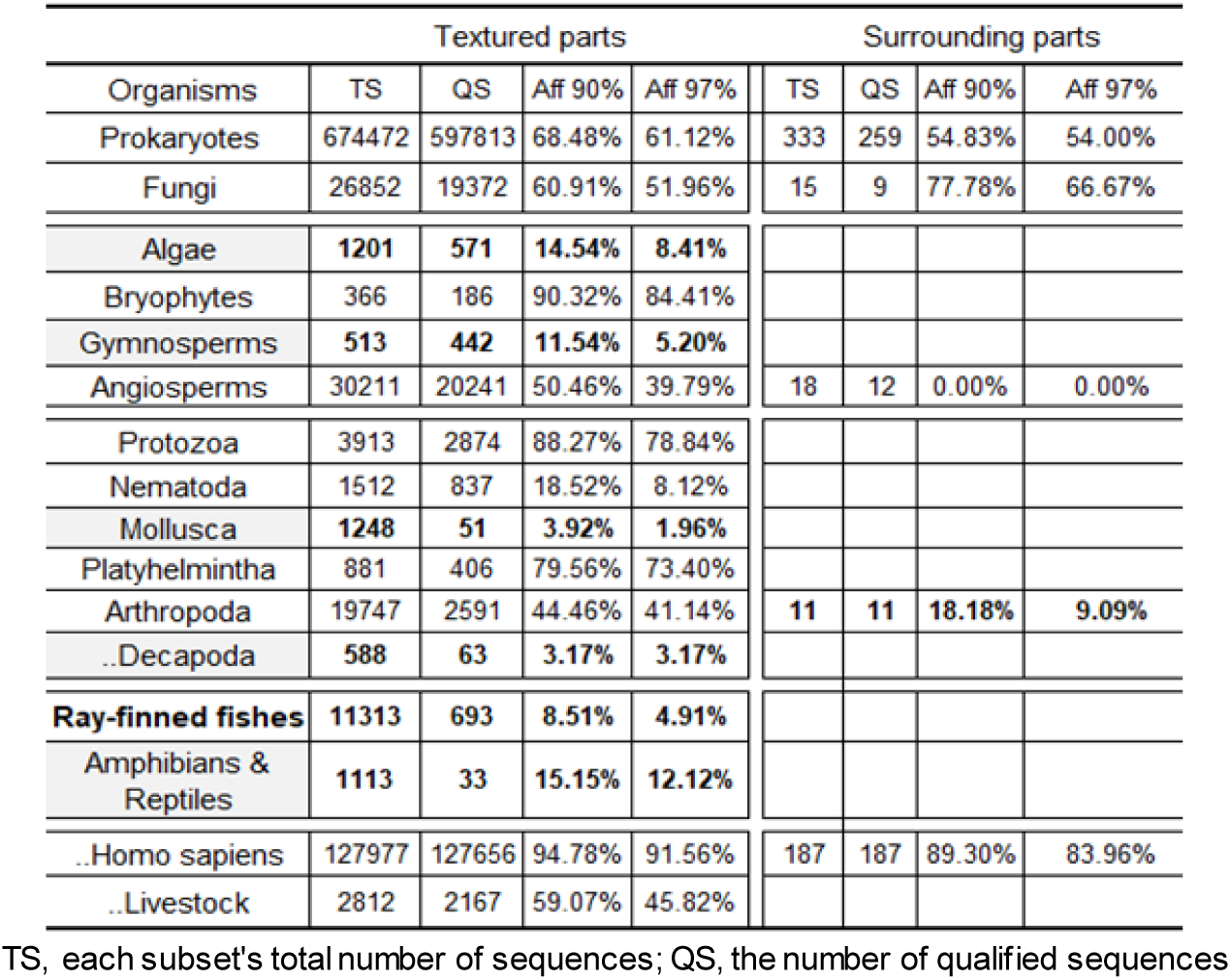
Sequencing reads and their matching lineage genomes.

| Organisms | Textured parts |  |  |  | Surrounding parts |  |  |  |
| --- | --- | --- | --- | --- | --- | --- | --- | --- |
|  | TS | QS | Aff 90% | Aff 97% | TS | QS | Aff 90% | Aff 97% |
| Prokaryotes | 674472 | 597813 | 68.48% | 61.12% | 333 | 259 | 54.83% | 54.00% |
| Fungi | 26852 | 19372 | 60.91% | 51.96% | 15 | 9 | 77.78% | 66.67% |
| Algae | 1201 | 571 | 14.54% | 8.41% |  |  |  |  |
| Bryophytes | 366 | 186 | 90.32% | 84.41% |  |  |  |  |
| Gymnosperms | 513 | 442 | 11.54% | 5.20% |  |  |  |  |
| Angiosperms | 30211 | 20241 | 50.46% | 39.79% | 18 | 12 | 0.00% | 0.00% |
| Protozoa | 3913 | 2874 | 88.27% | 78.84% |  |  |  |  |
| Nematoda | 1512 | 837 | 18.52% | 8.12% |  |  |  |  |
| Mollusca | 1248 | 51 | 3.92% | 1.96% |  |  |  |  |
| Platyhelmintha | 881 | 406 | 79.56% | 73.40% |  |  |  |  |
| Arthropoda | 19747 | 2591 | 44.46% | 41.14% | 11 | 11 | 18.18% | 9.09% |
| ..Decapoda | 588 | 63 | 3.17% | 3.17% |  |  |  |  |
| **Ray-finned fishes** | **11313** | **693** | **8.51%** | **4.91%** |  |  |  |  |
| Amphibians & Reptiles | 1113 | 33 | 15.15% | 12.12% |  |  |  |  |
| ..Homo sapiens | 127977 | 127656 | 94.78% | 91.56% | 187 | 187 | 89.30% | 83.96% |
| ..Livestock | 2812 | 2167 | 59.07% | 45.82% |  |  |  |  |

TS, each subset's total number of sequences; QS, the number of qualified sequences

**Table S2.** Some characteristics of DNA sequences in the ray-finned fish subset.

| Markers | *Homo sapiens* |  | oriDNA |  |
| --- | --- | --- | --- | --- |
|  | Mean | SD | Mean | SD |
| Length | 135.65 | 23.8 | 139 | 21.28 |
| Per Identity | 98.32% | 6.46% | 68.65% | 21.05% |
| Query cover | 98.82% | 6.08% | 77.02% | 23.37% |
| Subset Affinity | 97.15% | 0.39% | 52.87% | 4.92% |
| Gap(s)/Sequence | 0.00 | 0.27 | 2.25 | 2.70 |
| GC% | 42.05% | 9.73% | 44.68% | 11.00% |
| QS/TS (%) | 99.70% |  | 6.14% |  |

**Table S3A.**
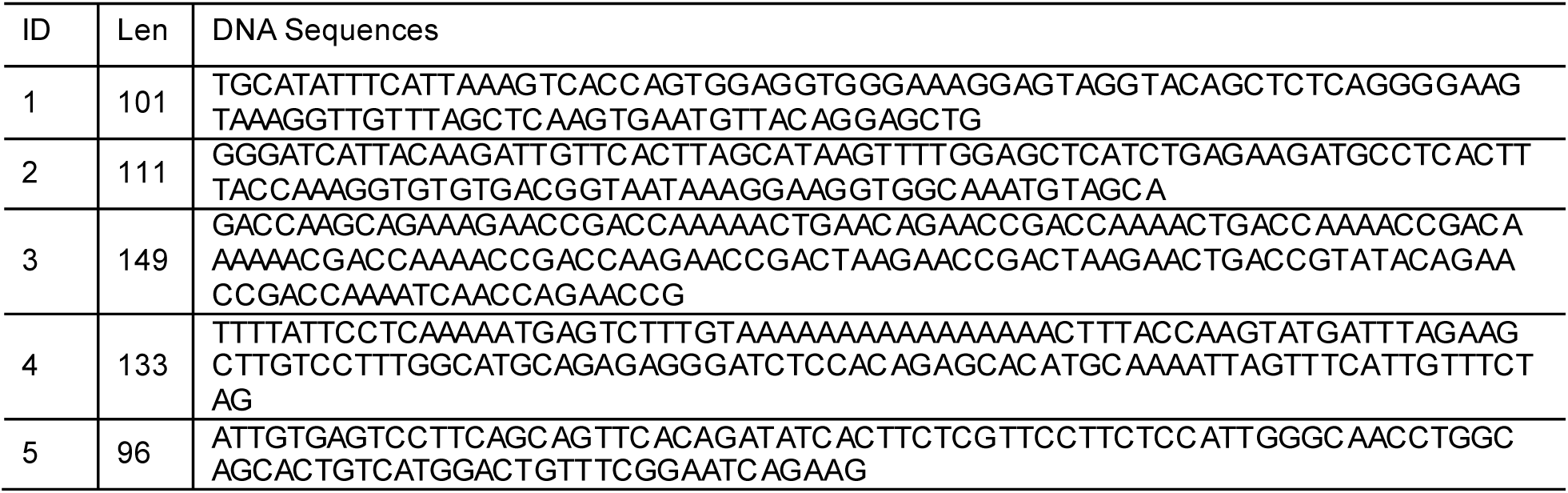
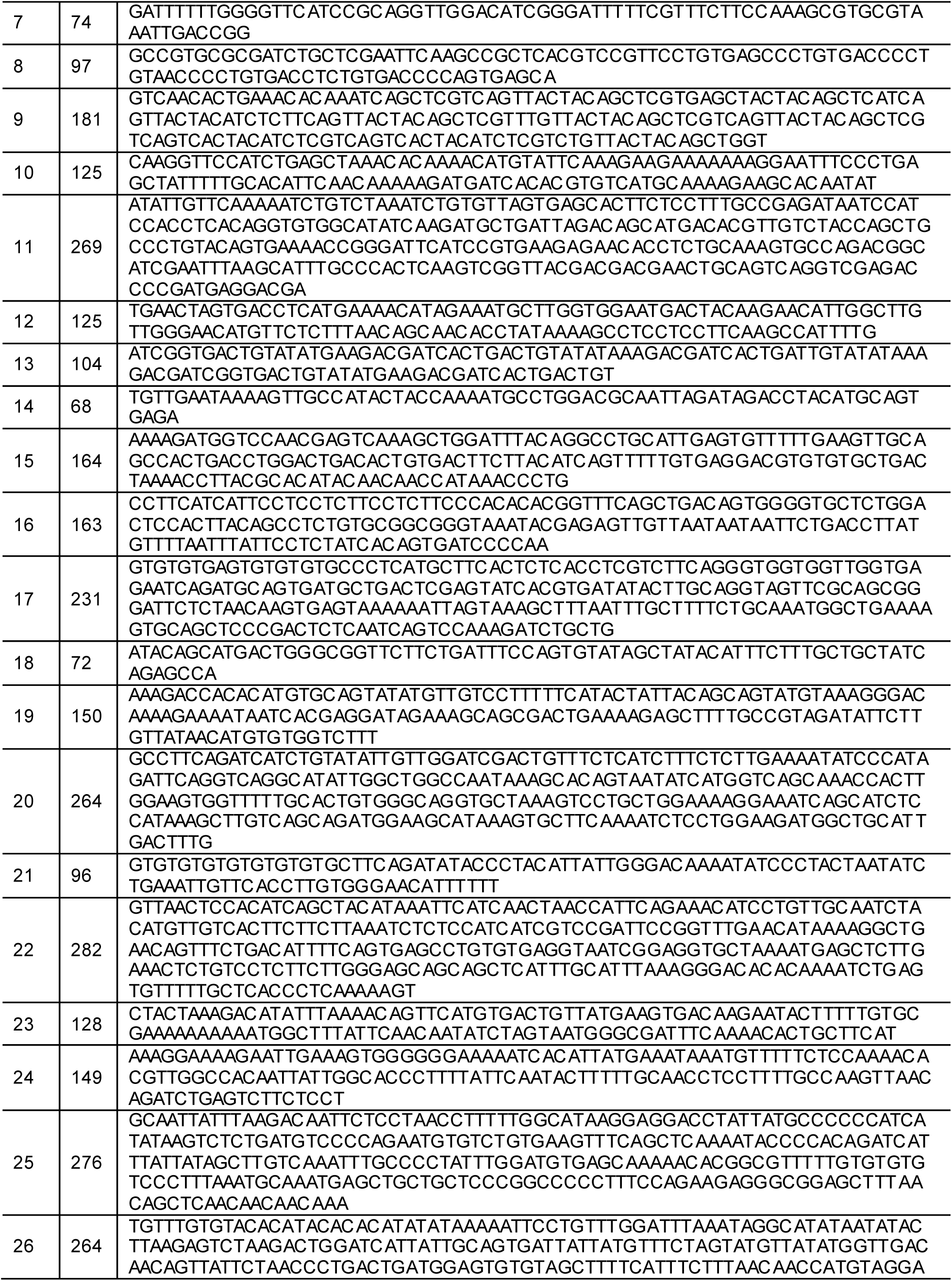
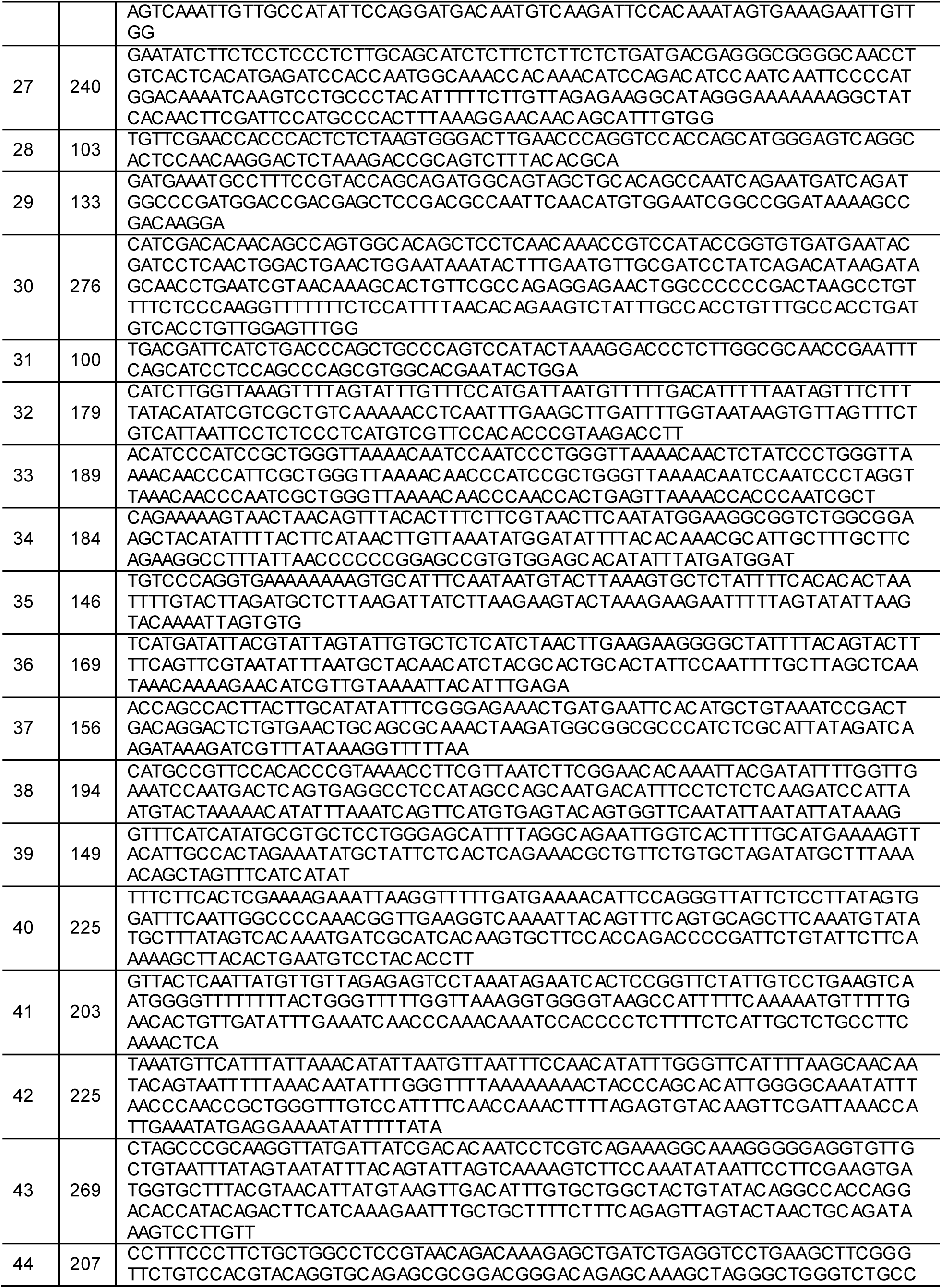
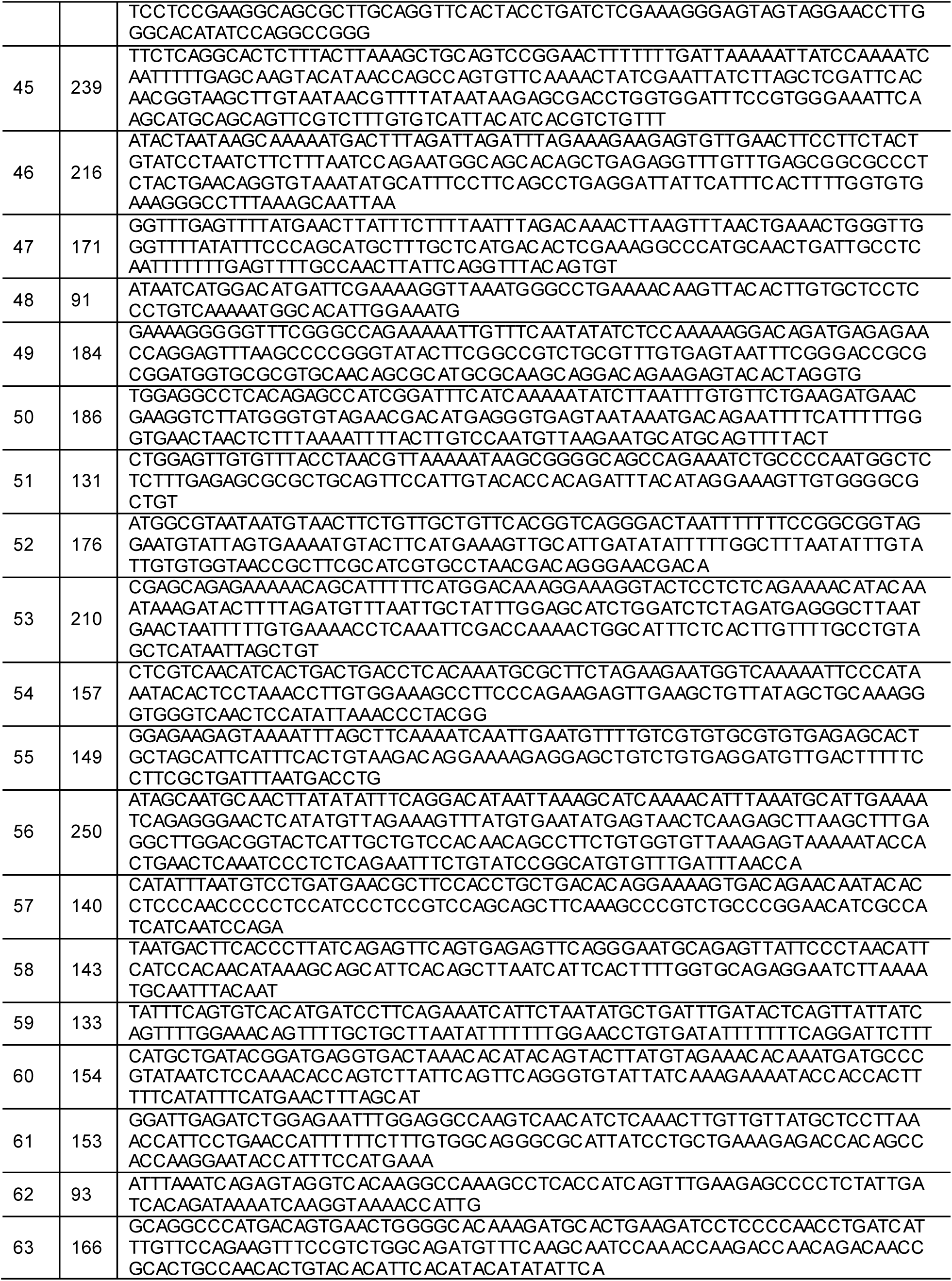
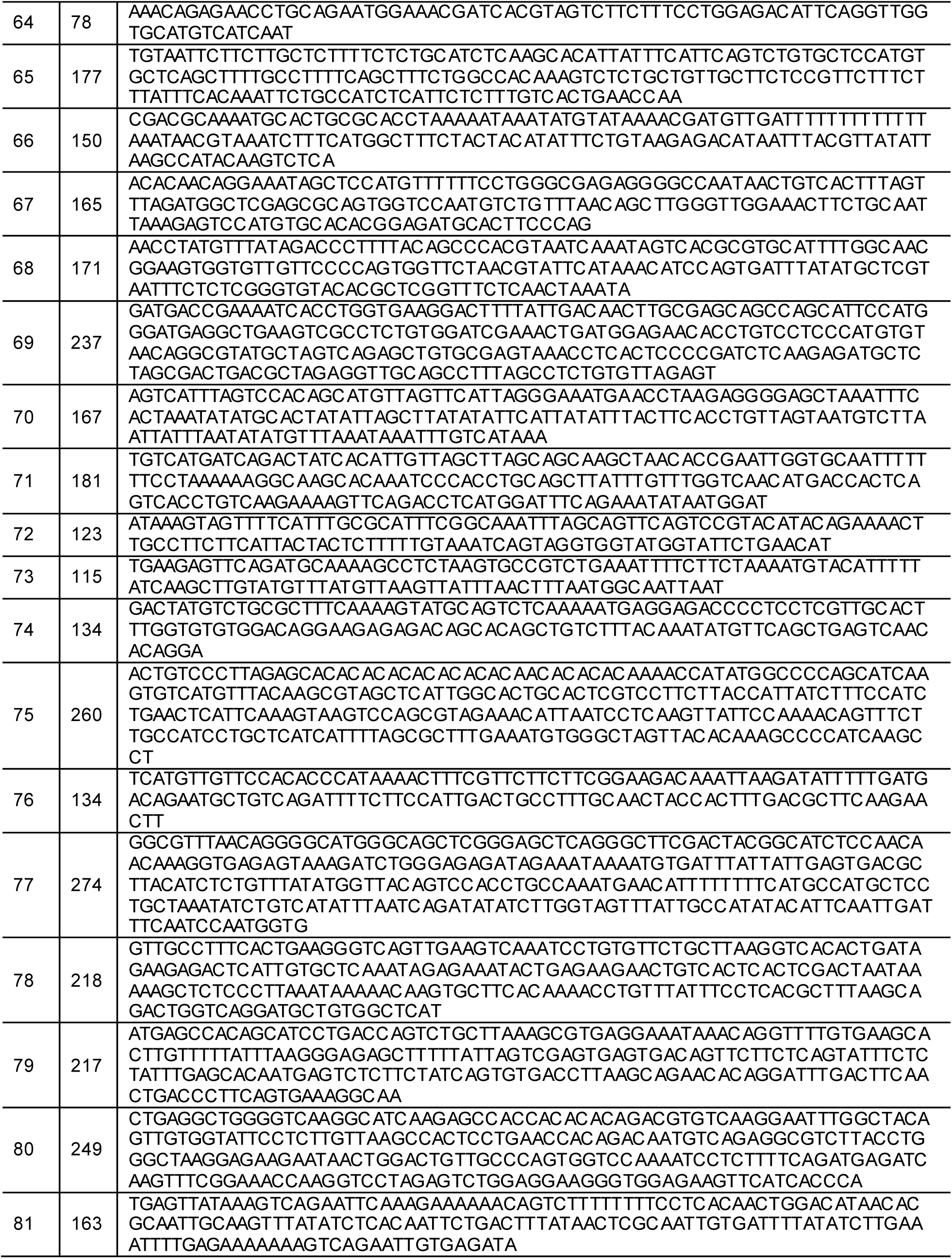
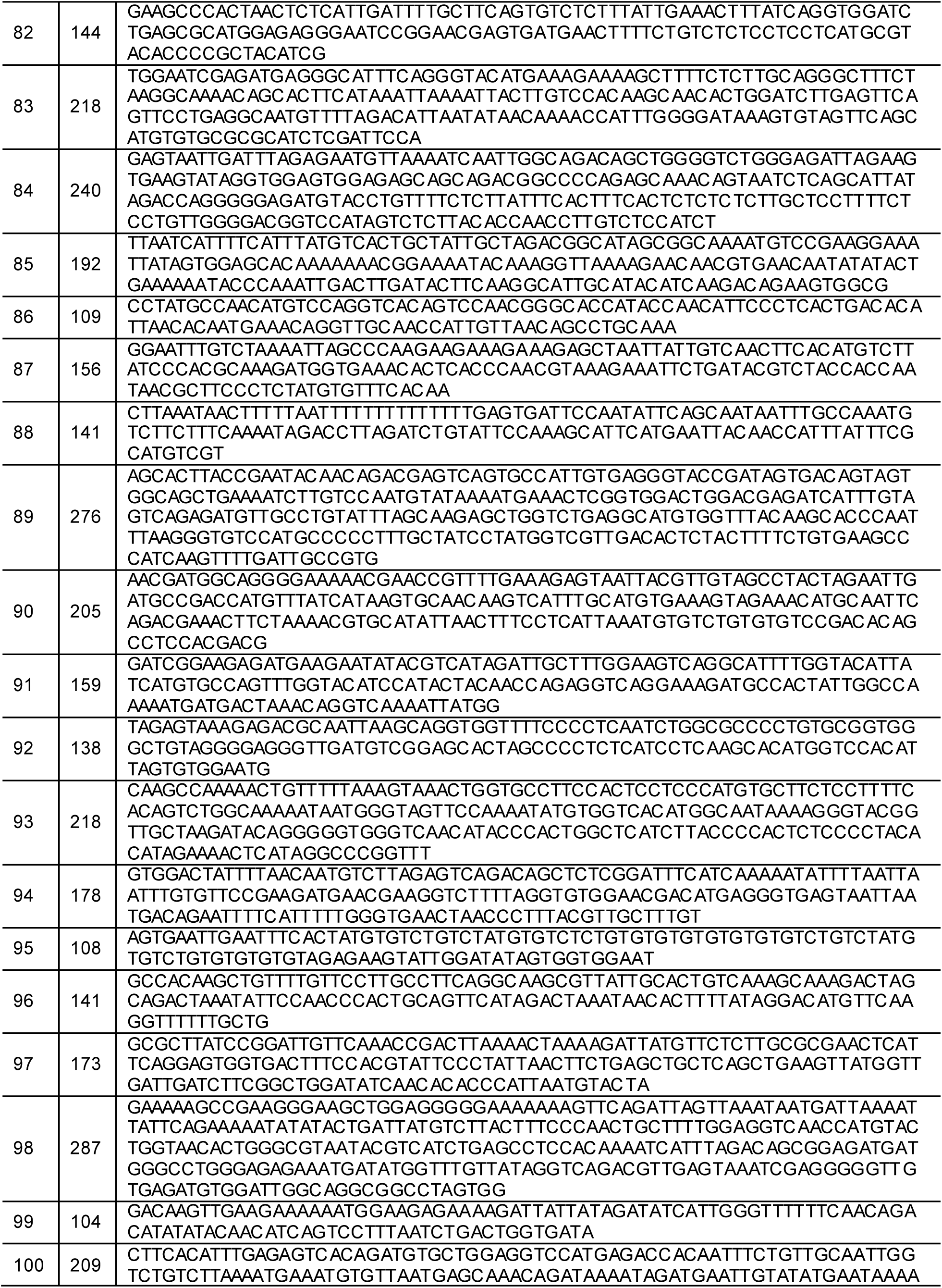
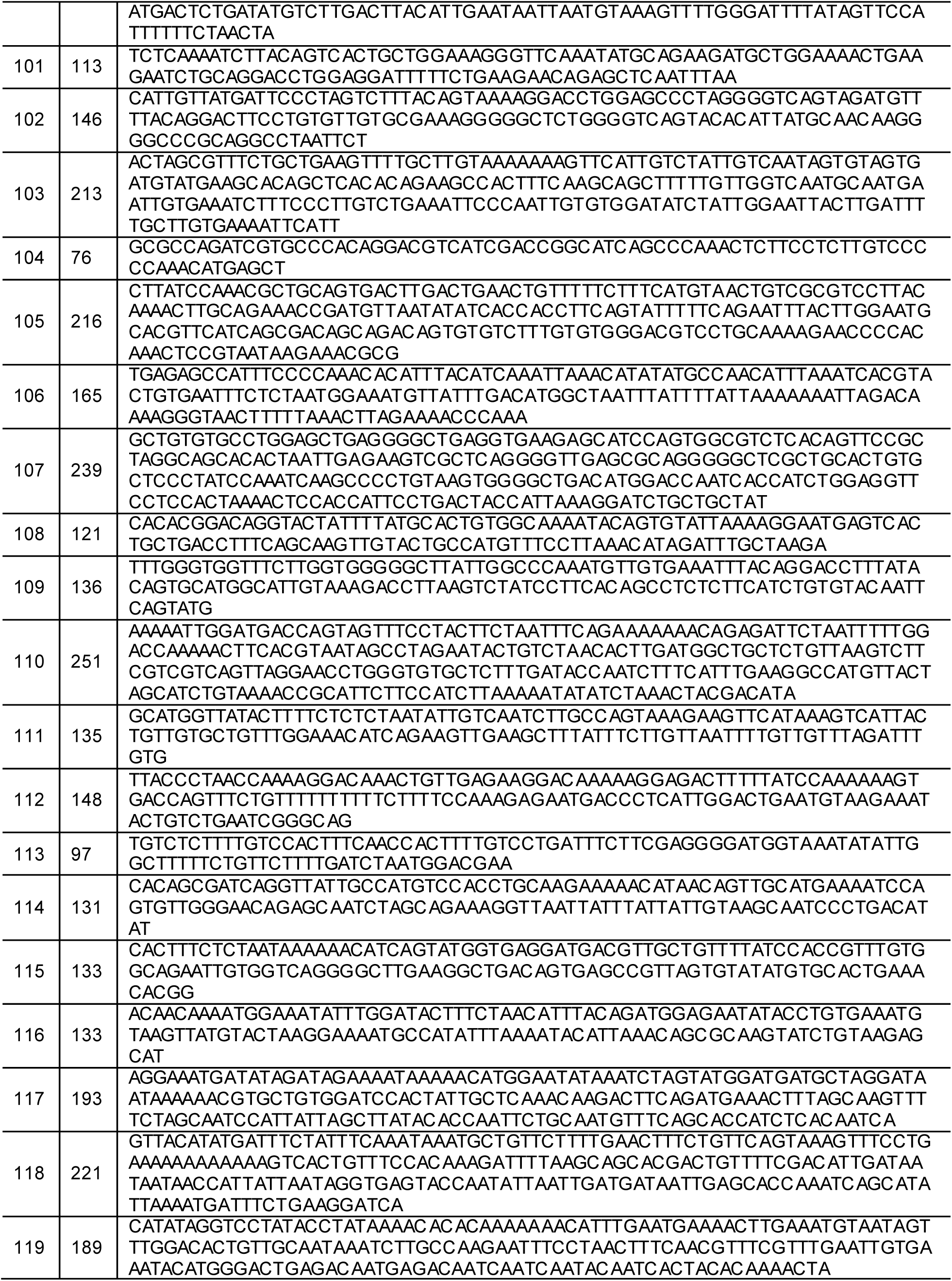
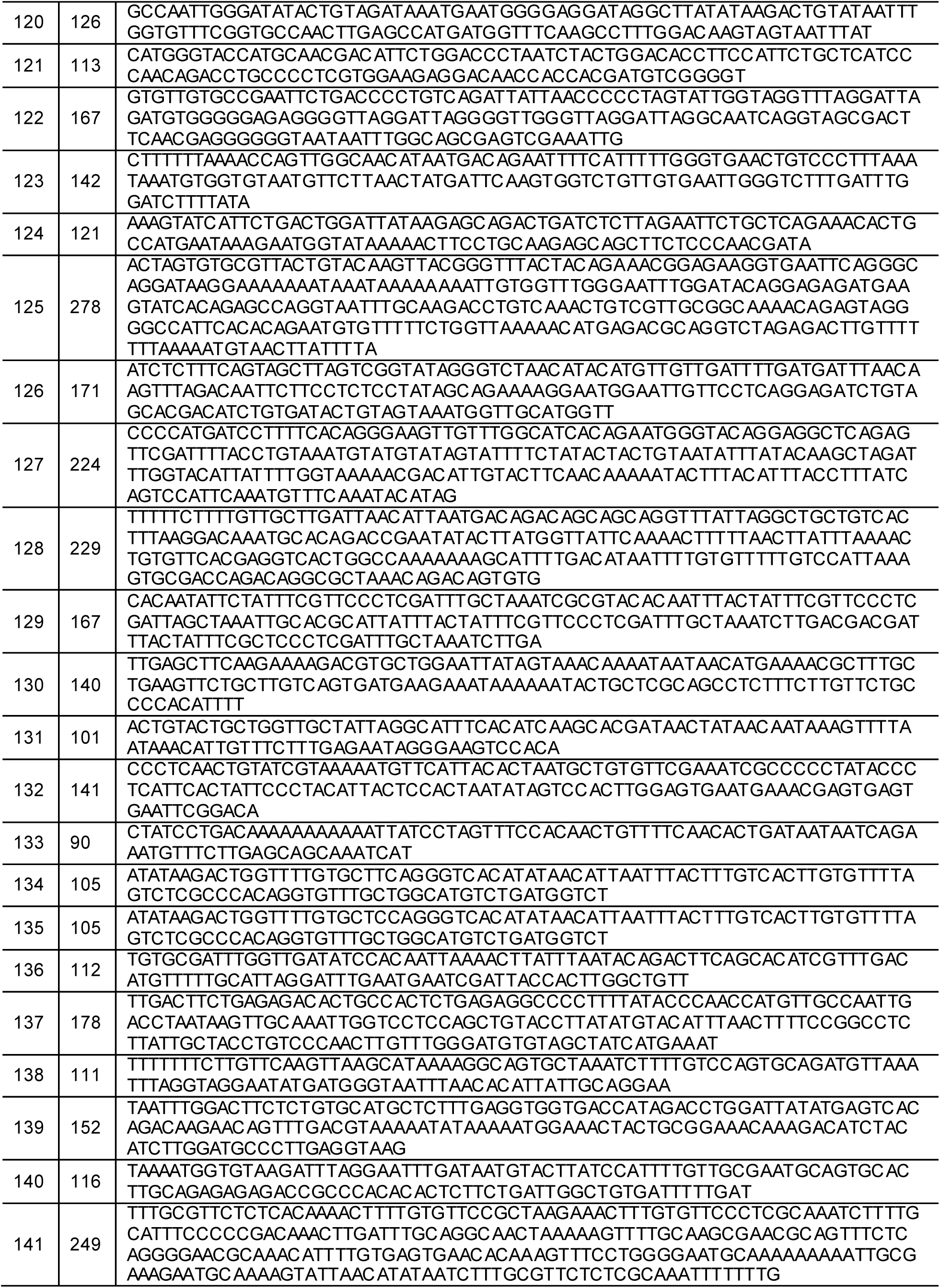
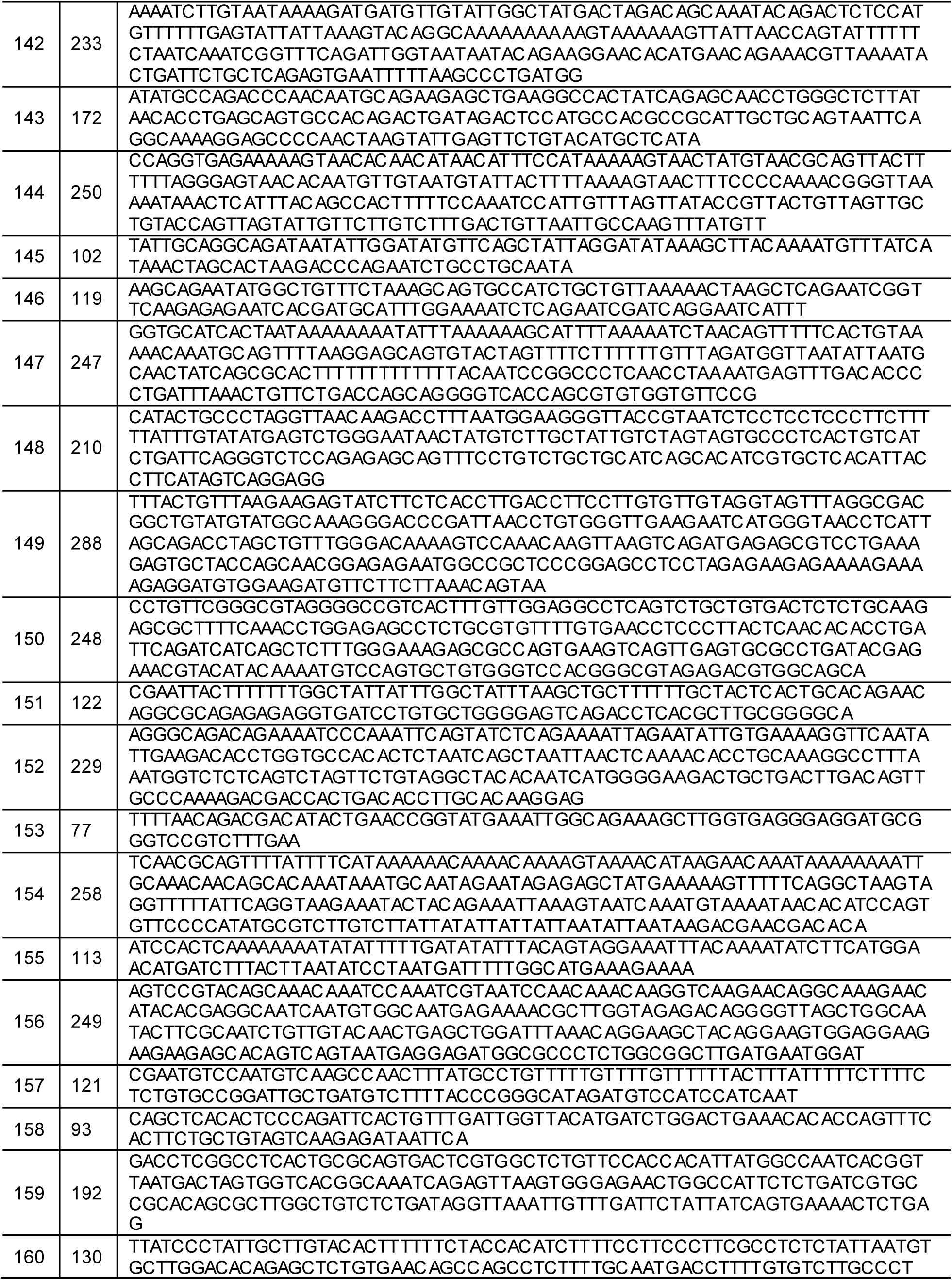
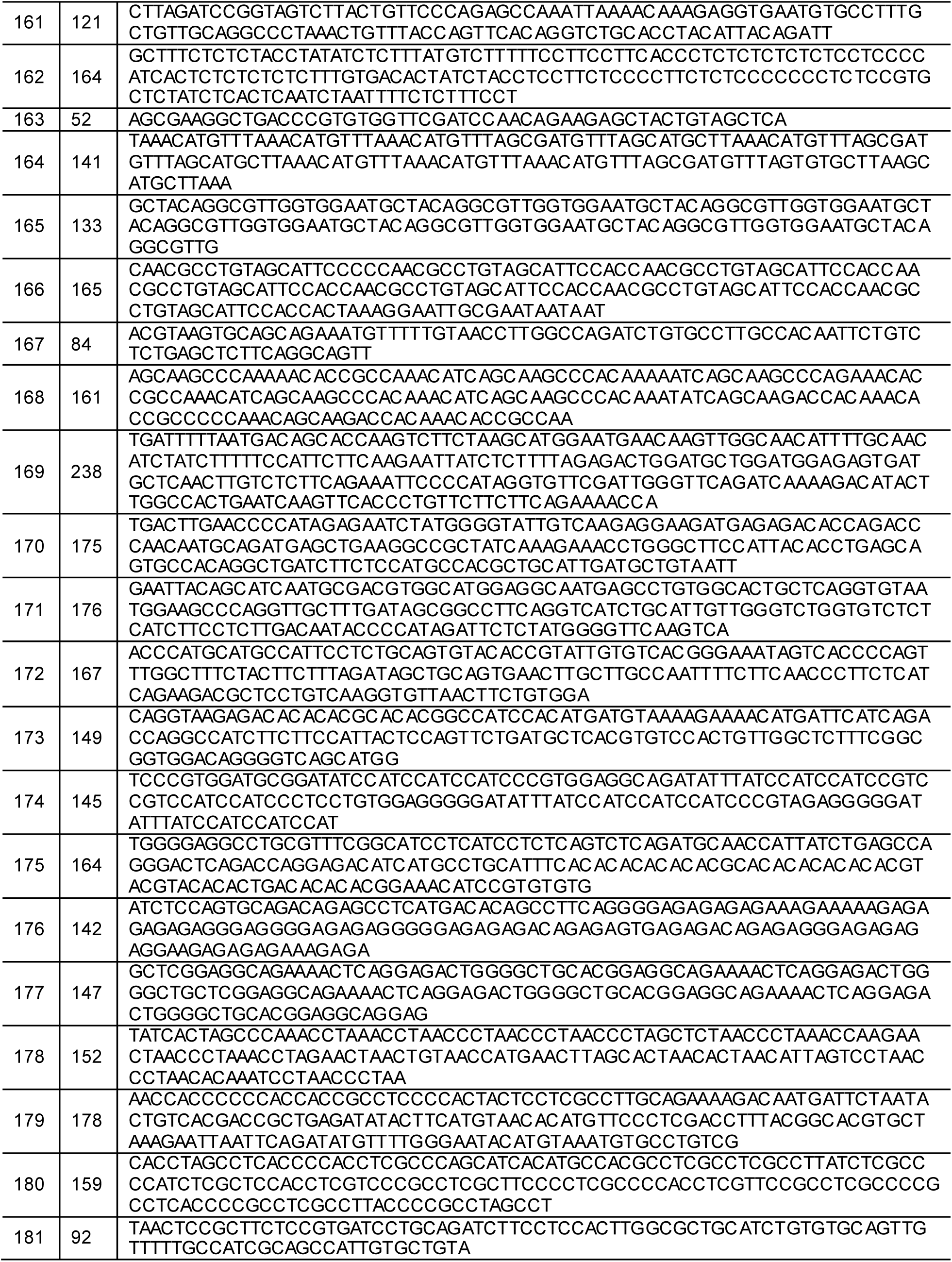
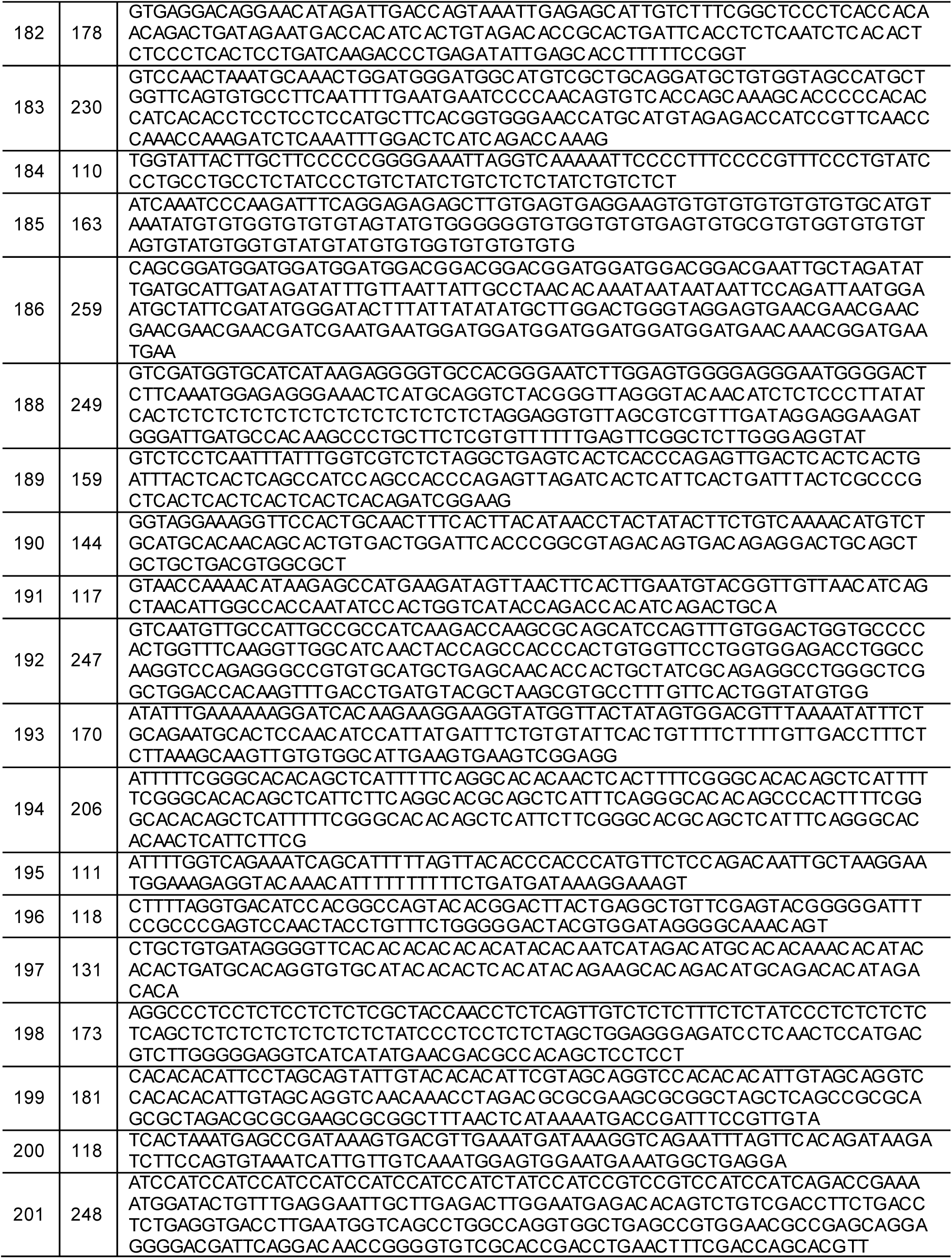
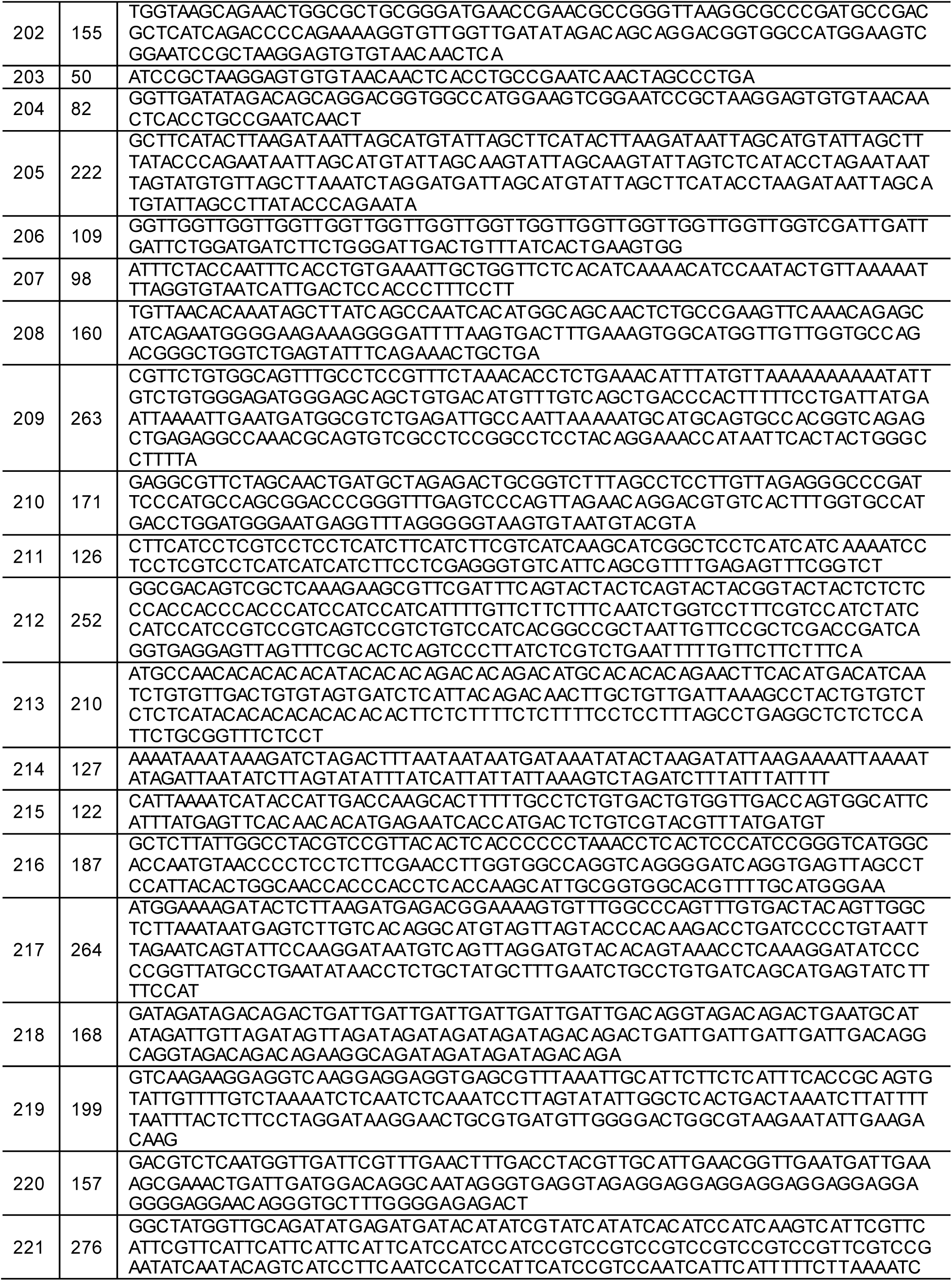
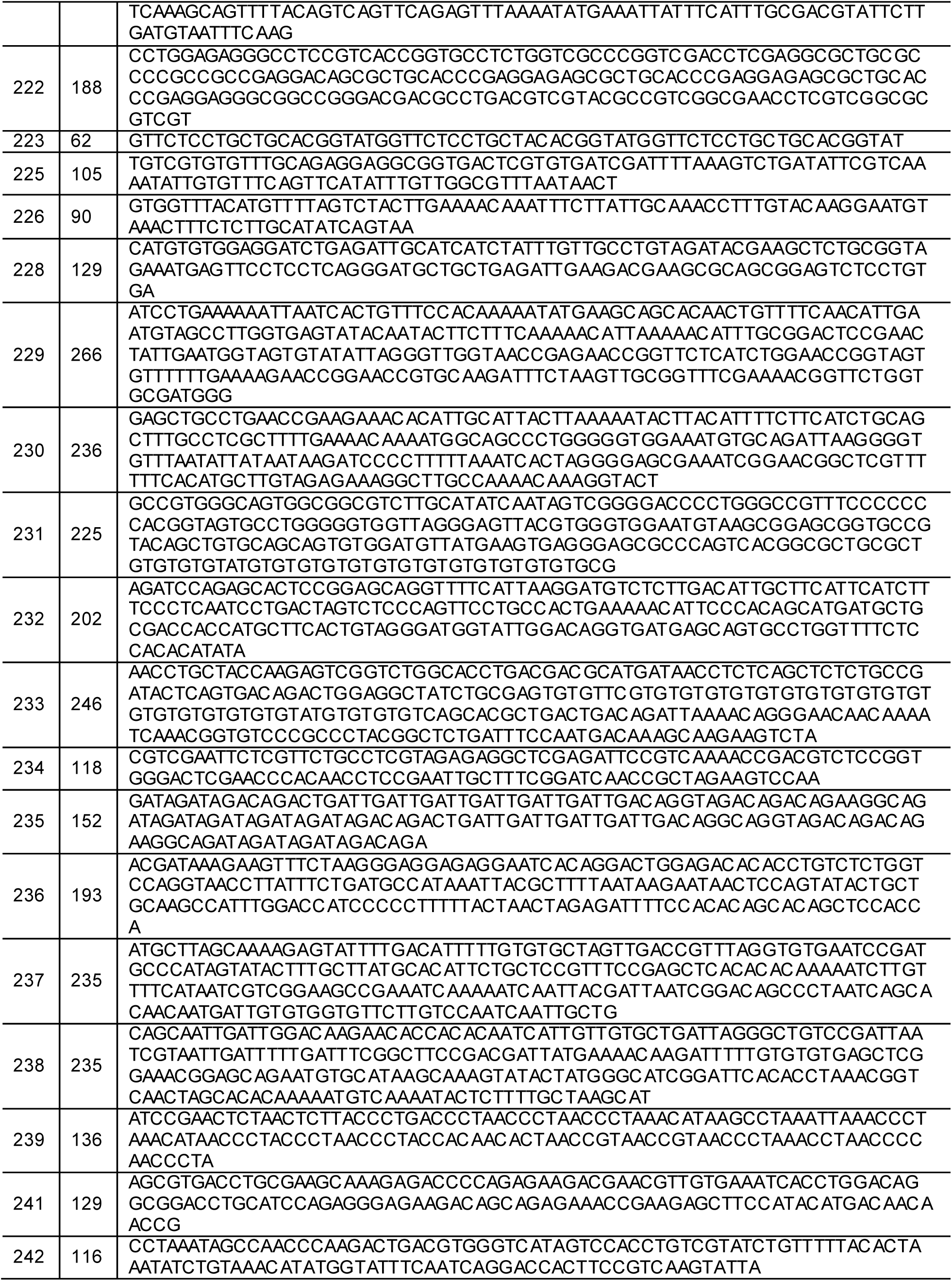
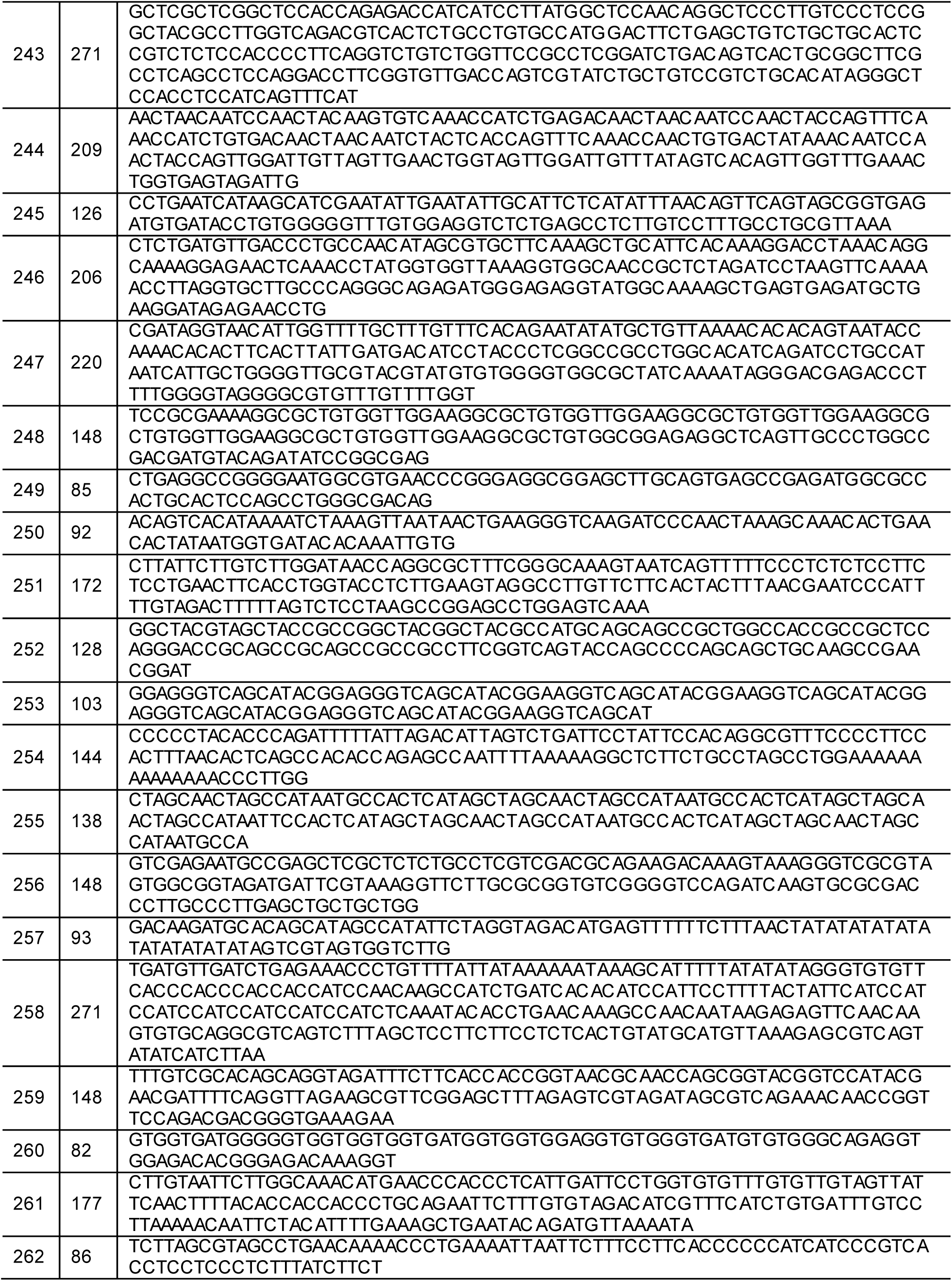
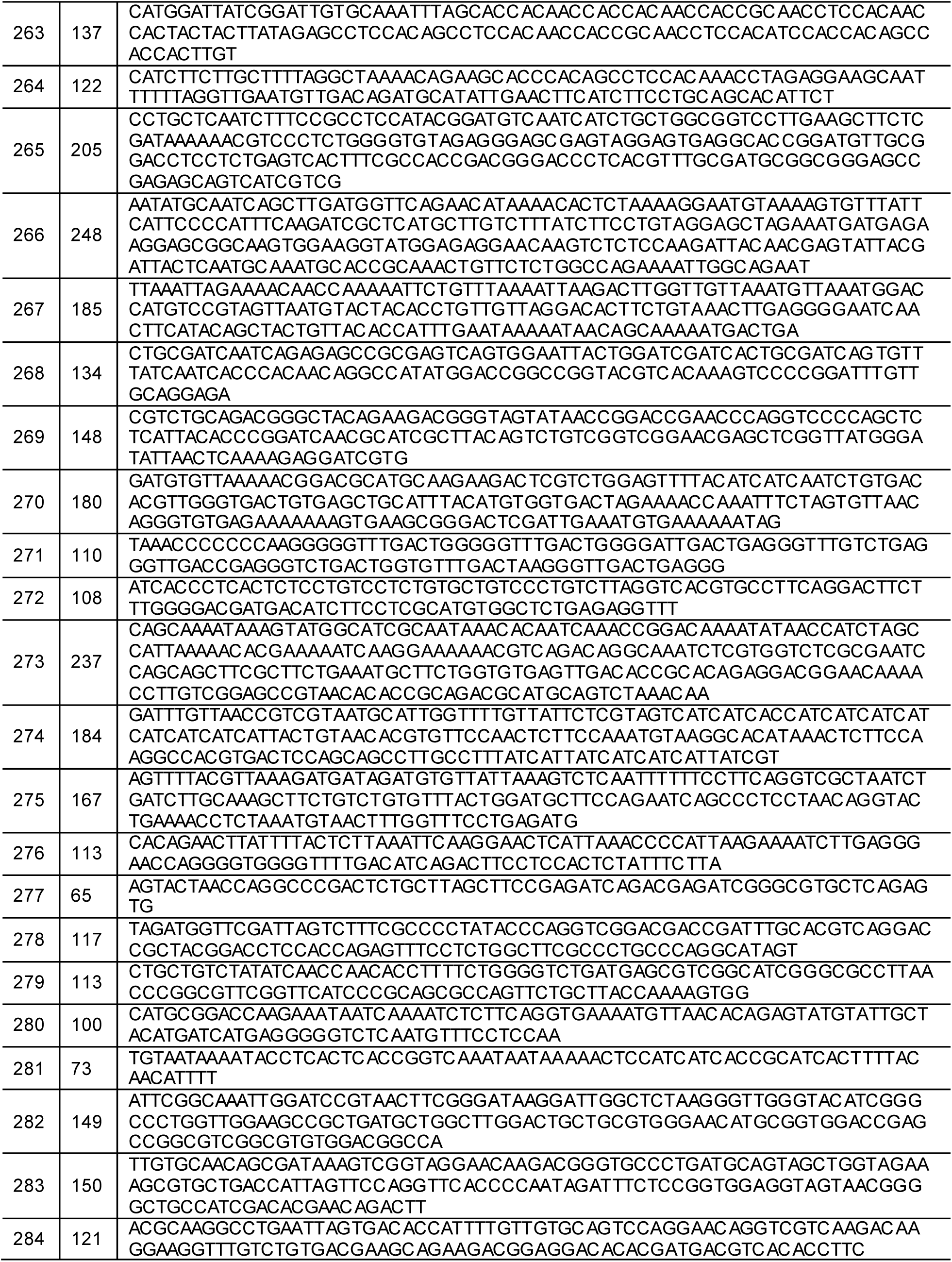
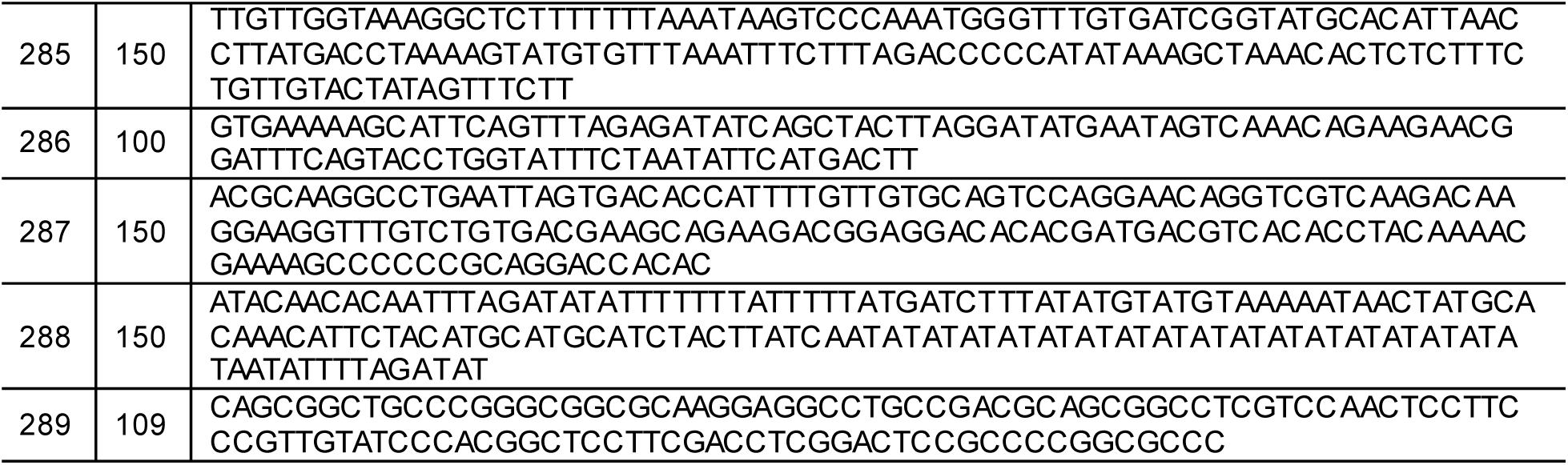
DNA sequences of the ray-finned fishes and the food web symbiotic species.

**Table S3B.**
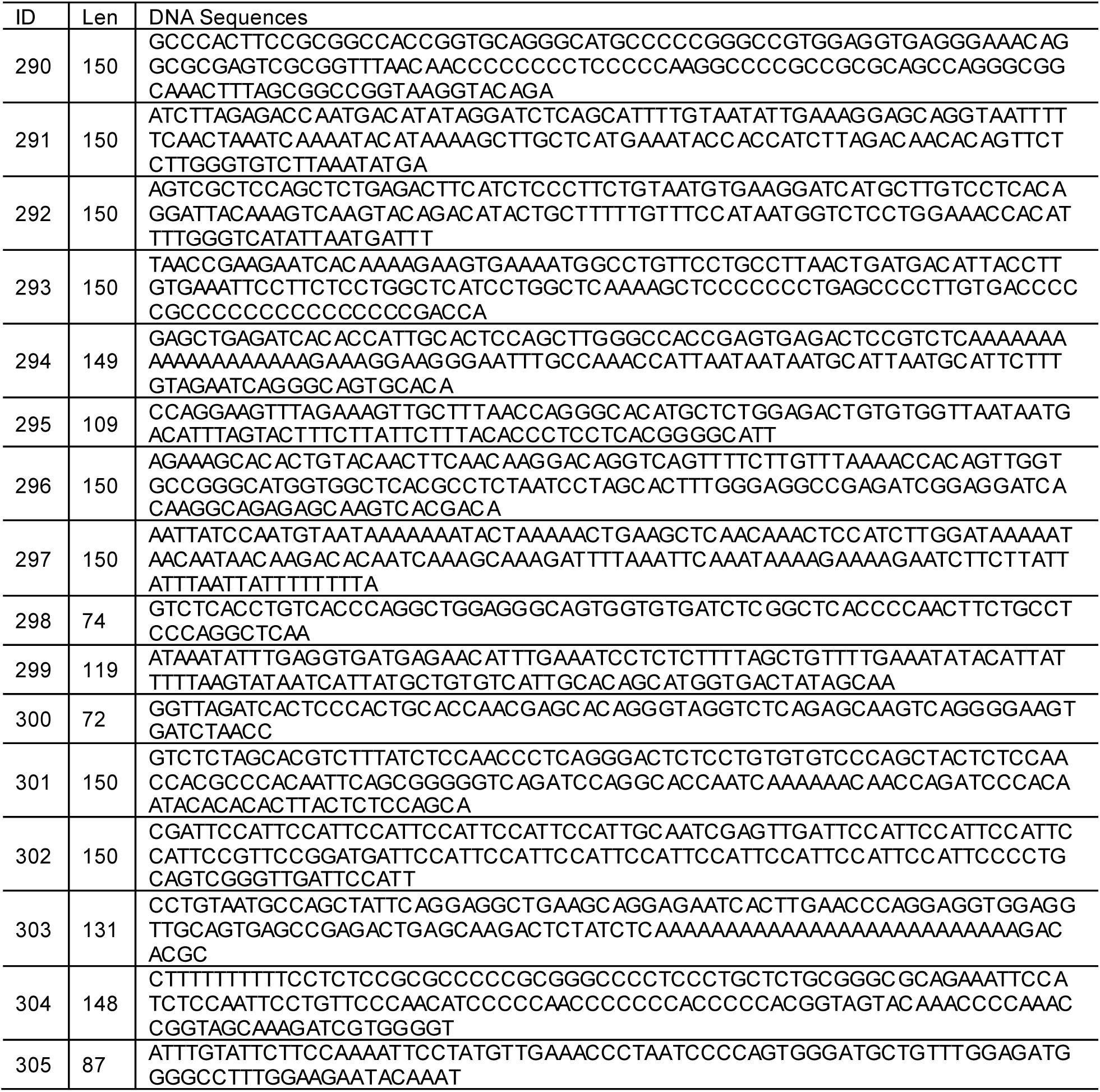
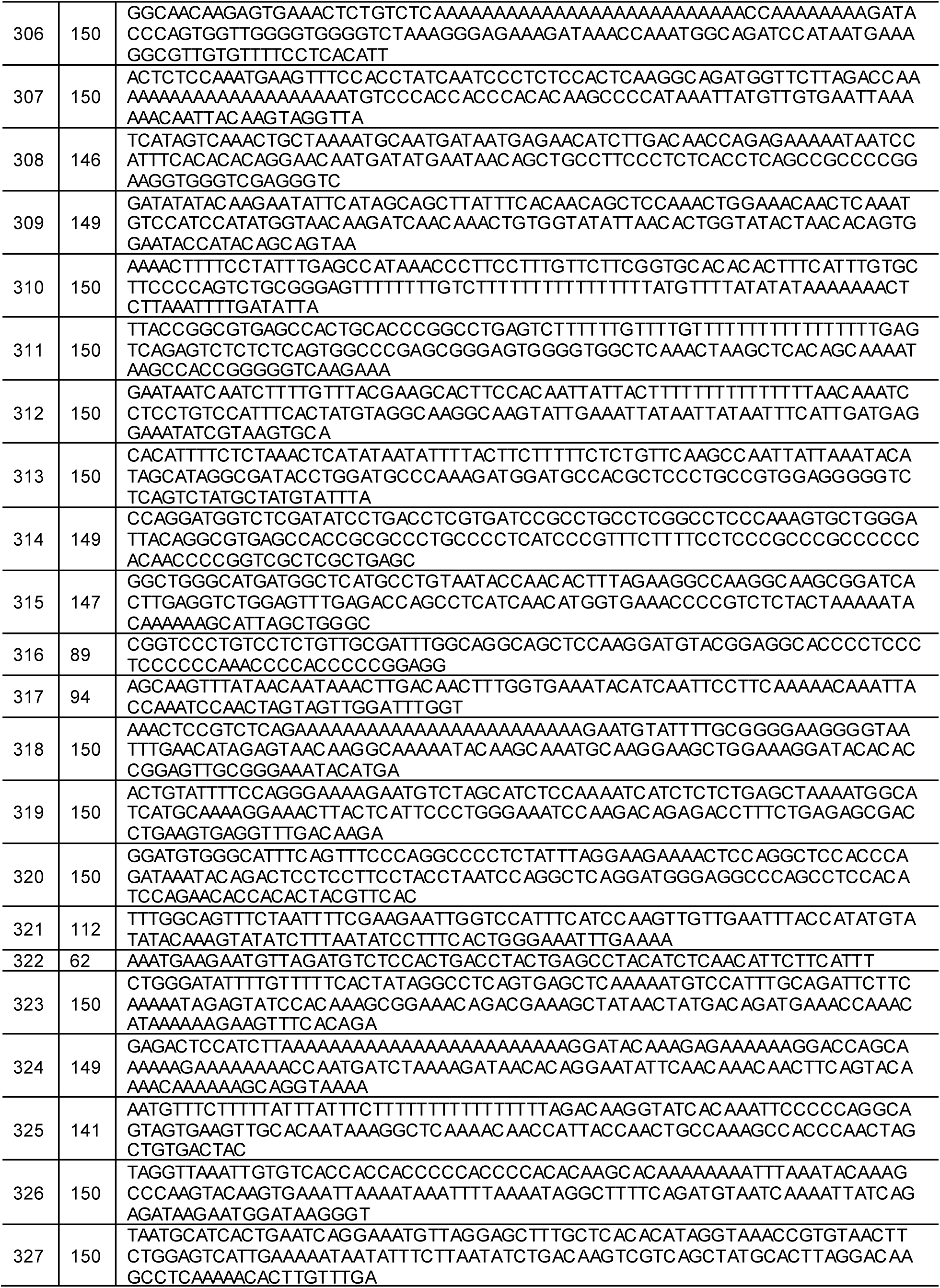
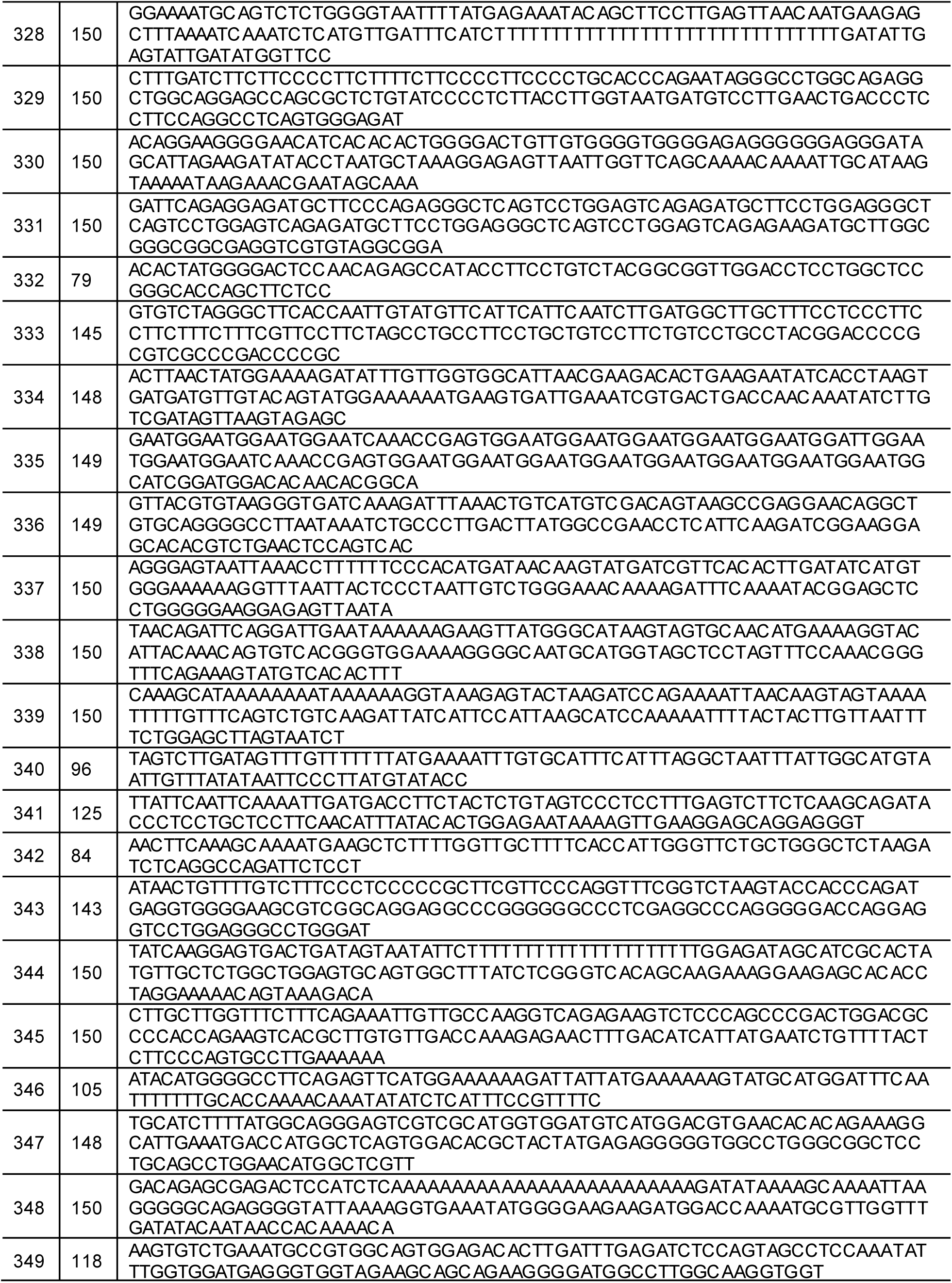
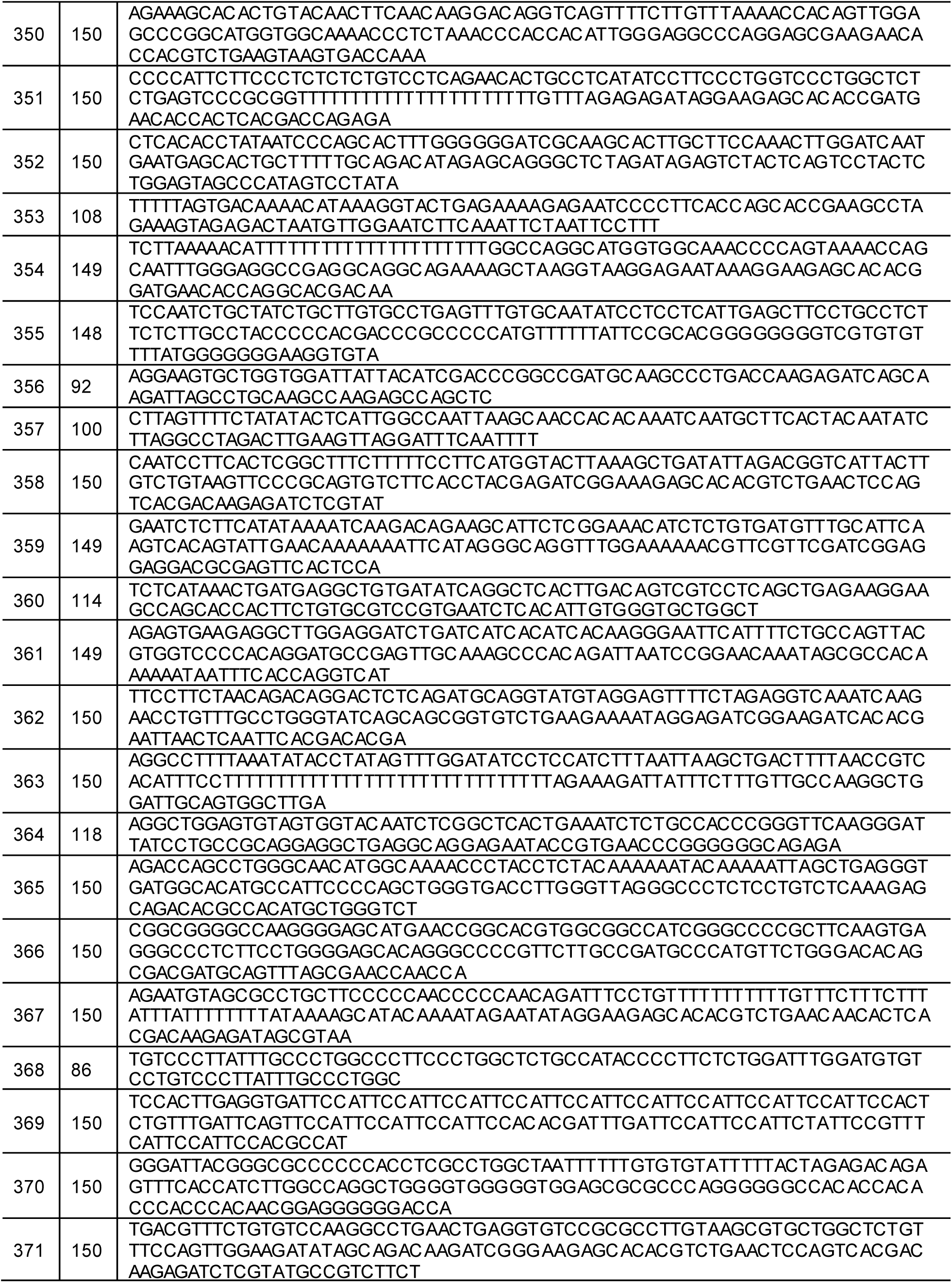
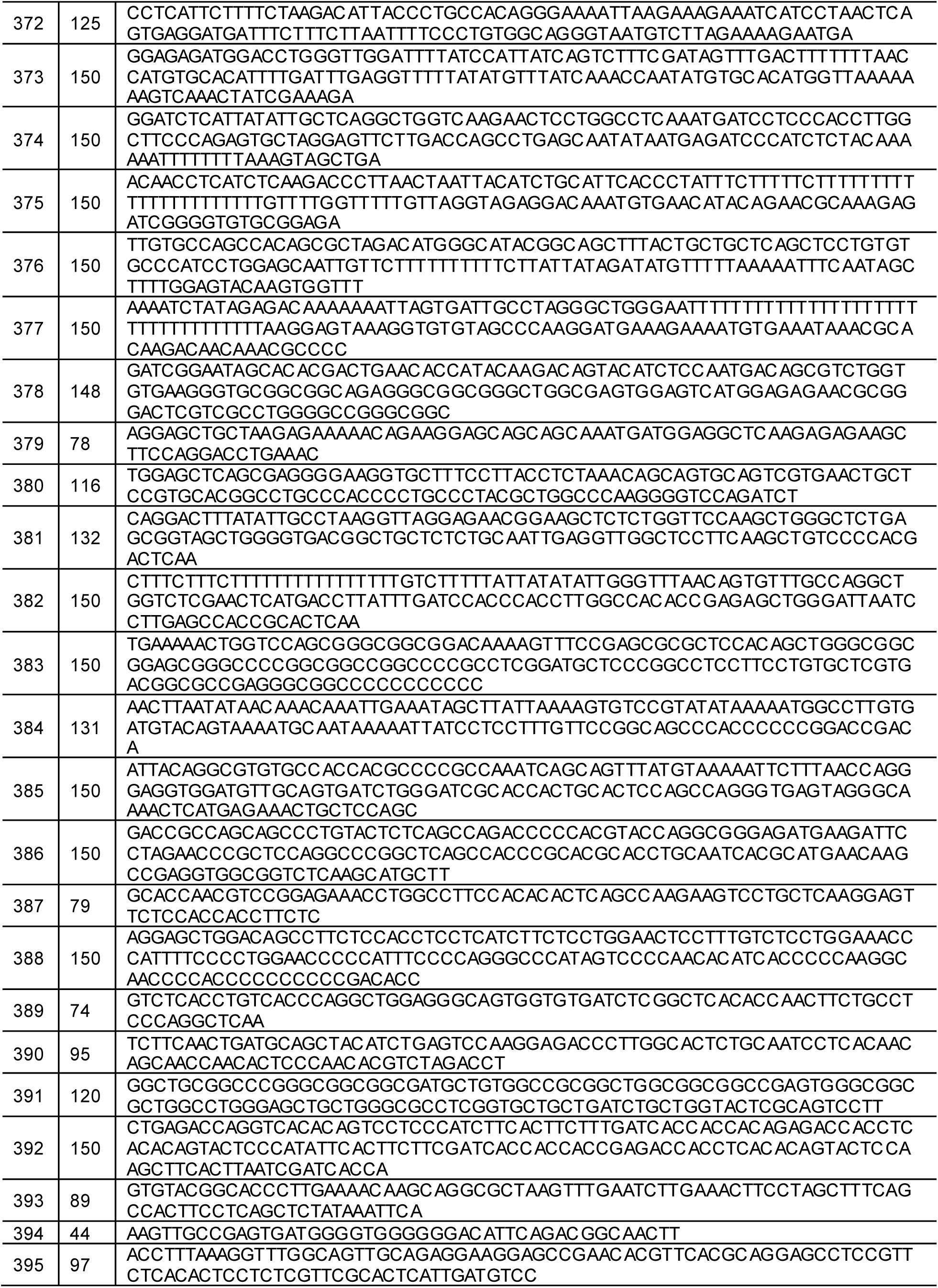
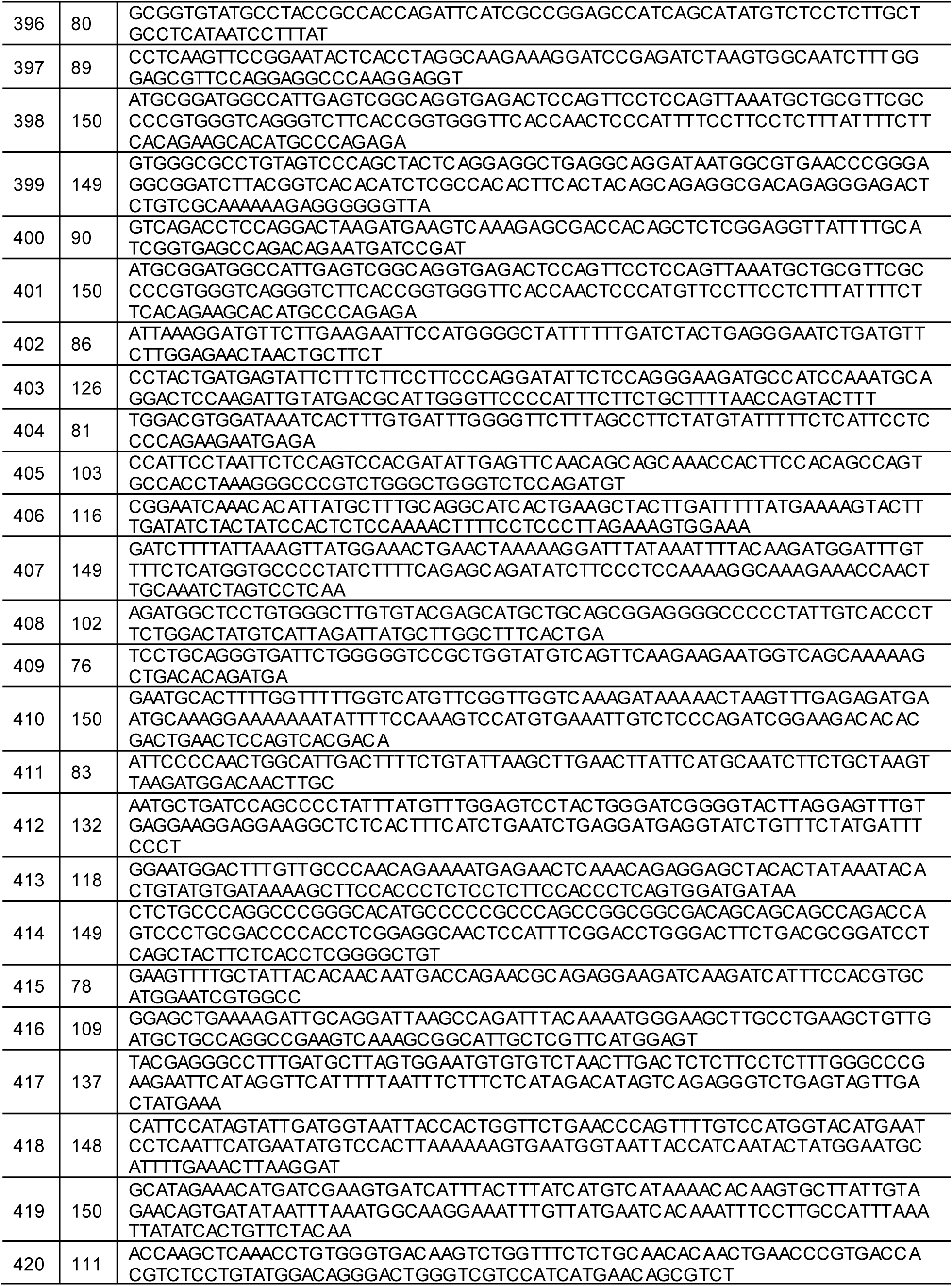
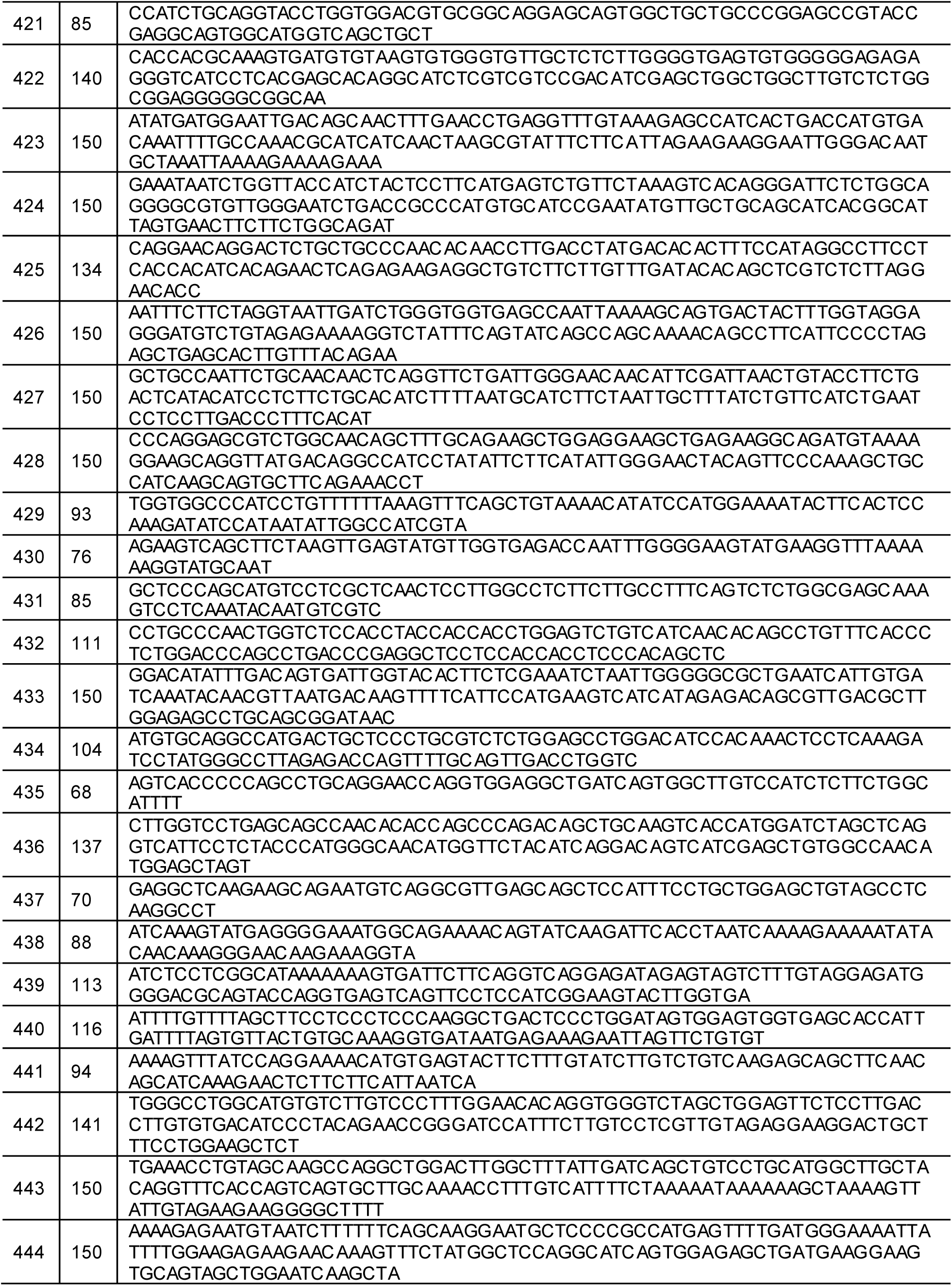
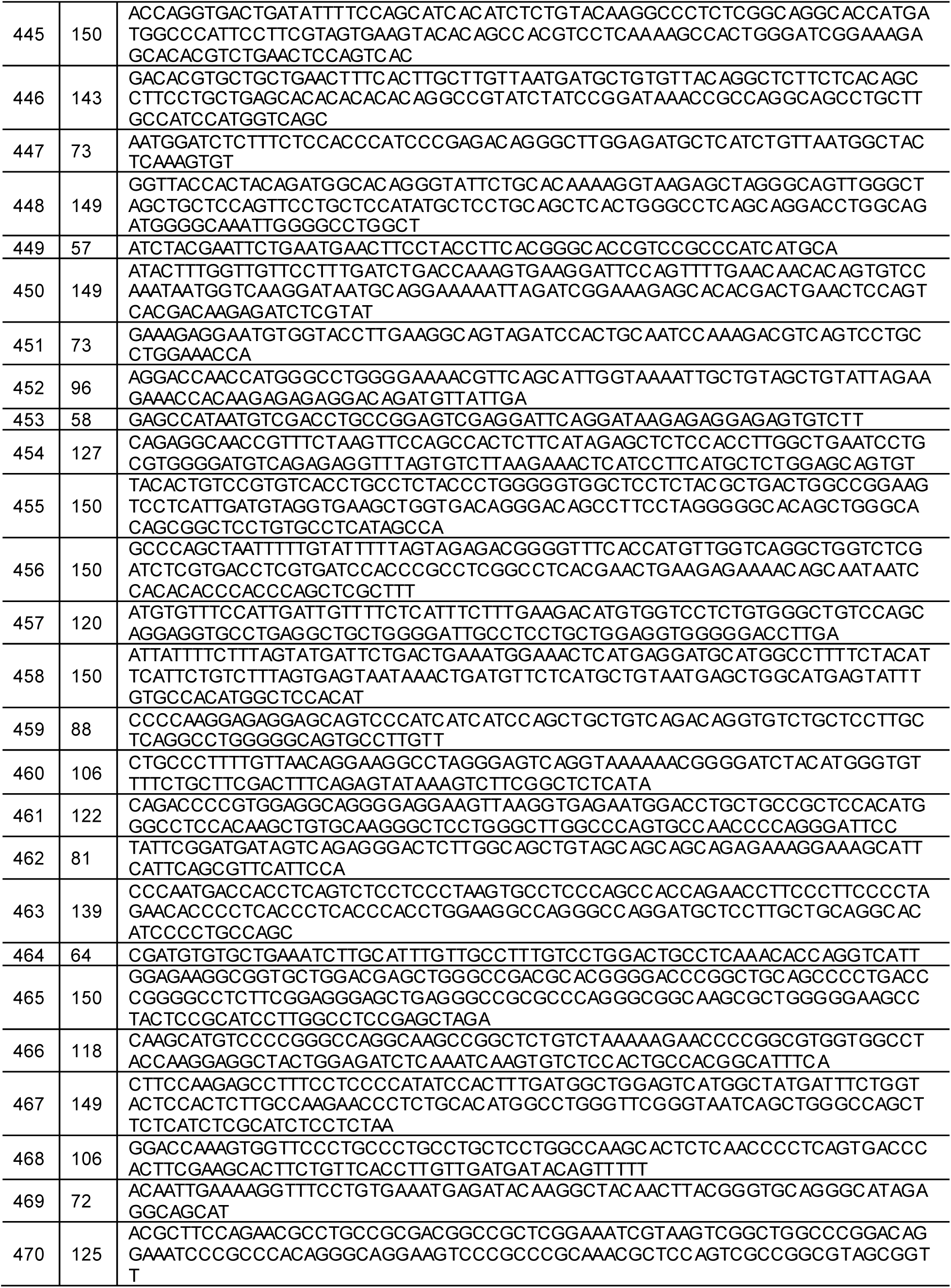
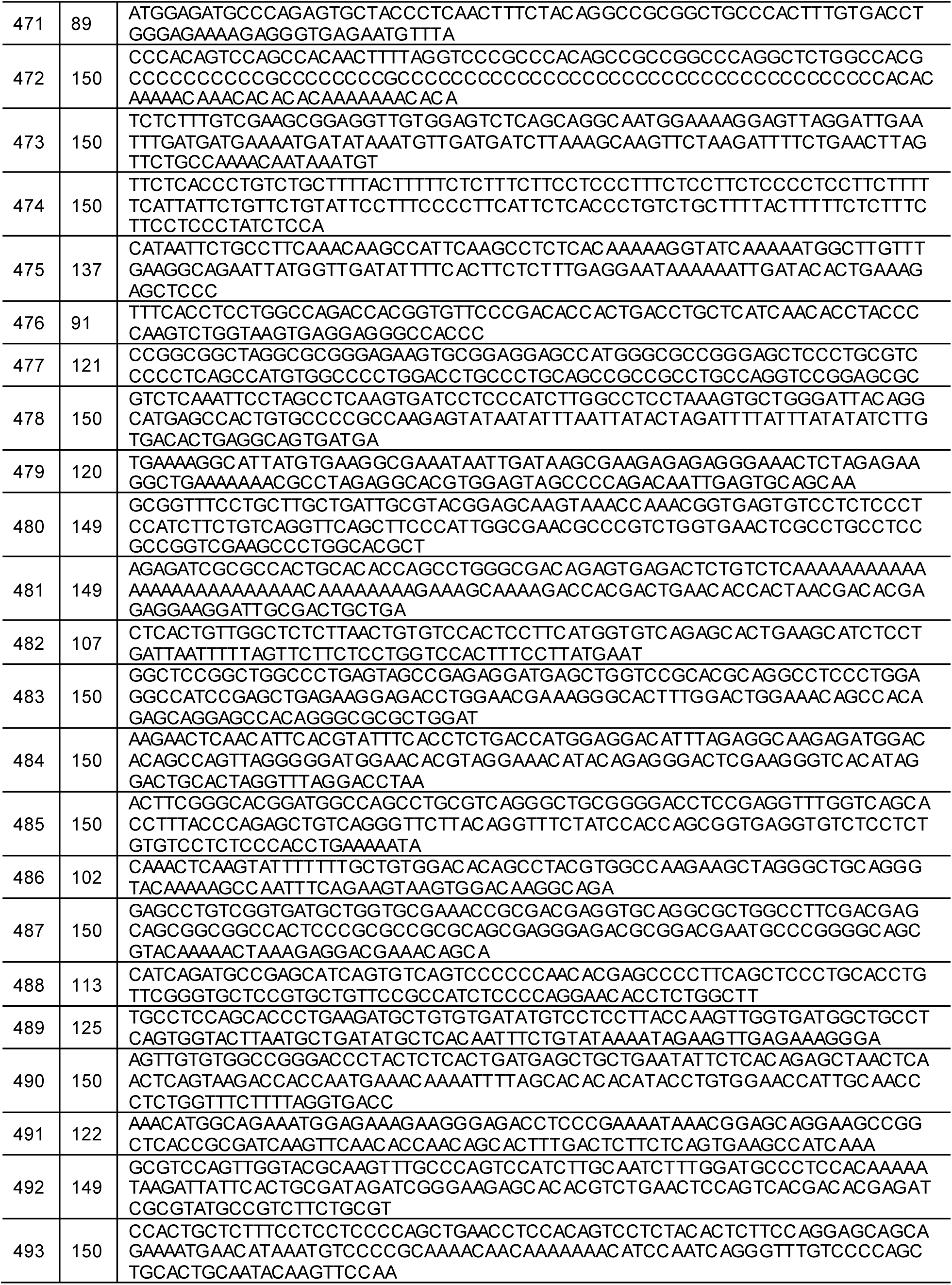
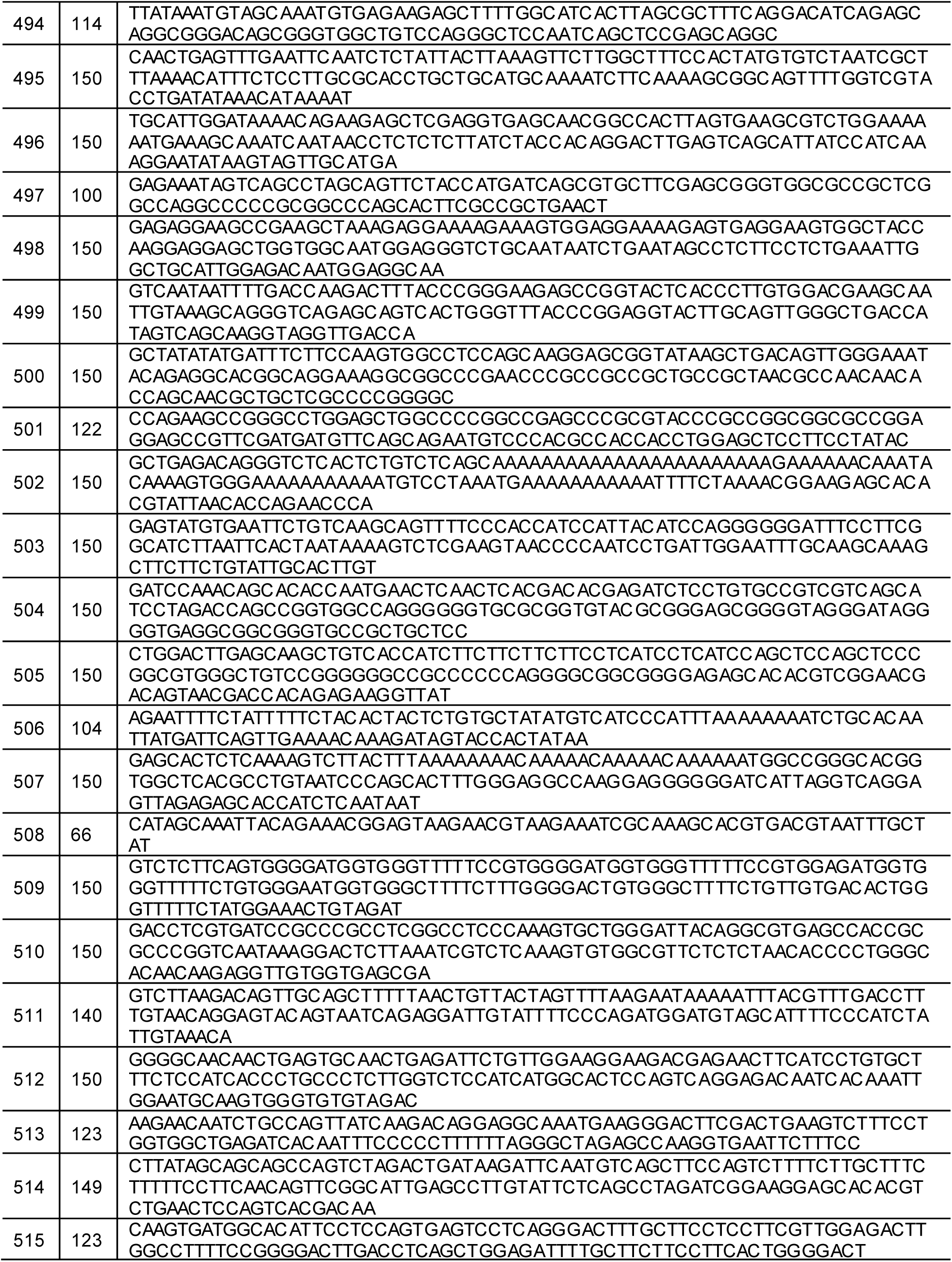
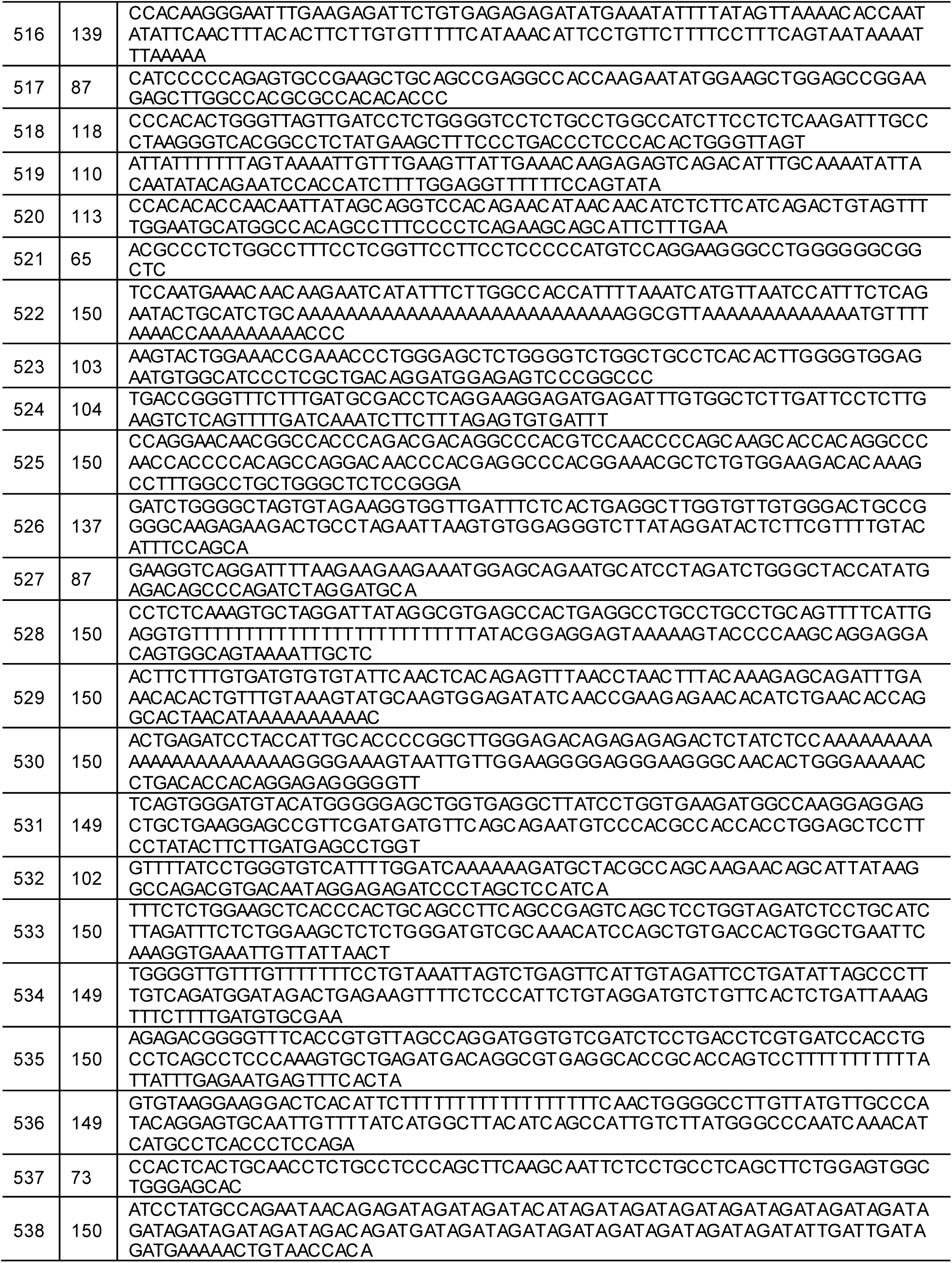
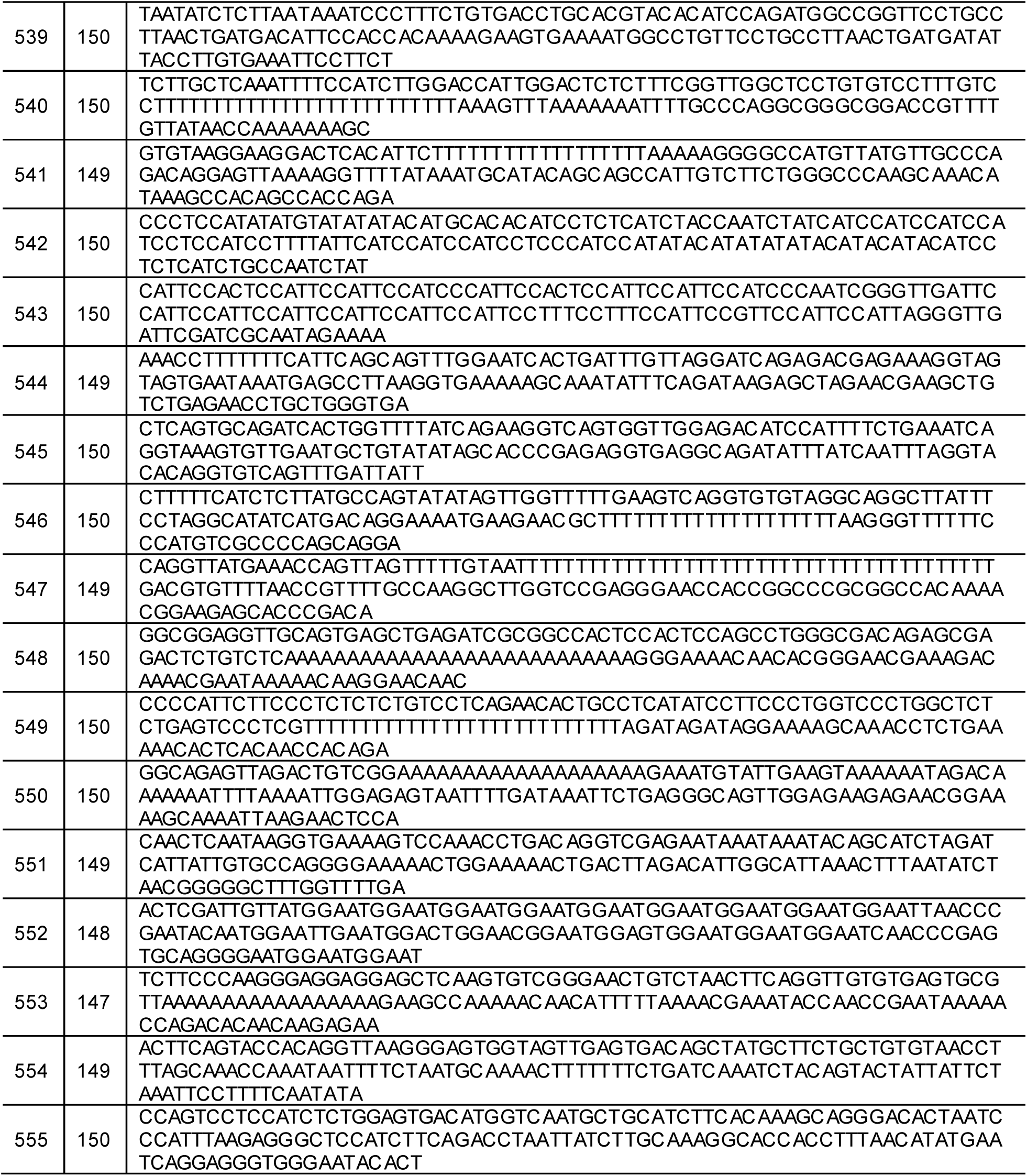
DNA fragments matching NHP’s sequences.

**Table S4.** Geographical classification of DNA sequences from the subset of ray-finned fishes.

| Group | Sequence ID | Number | Average affinity | E-value | Blast Mode |
| --- | --- | --- | --- | --- | --- |
| Non-native | #001 #002 #003 #007 #009 #010 #012 #013 #014 #015 #016 #164 #165 #166 #168 #<u>169</u> #017 #049 #176 #179 #180 #181 #182 #183 #190 #191 #192 #193 #194 #195 #196 #199 #205 #208 #223 #225 #226 #233 #236 #240 #246 #253 #254 #258 #259 #260 #261 #262 #272 | 49 | 80.34% | $\leq 4.00\text{E-}9$ | the MS mode |
| Native | #004 #<u>011</u> #018 #019 #<u>020</u> #021 #022 #023 #024 #025 #026 #207 #028 #029 #030 #031 #032 #033 #034 #035 #036 #037 #038 #039 #040 #041 #042 #043 #044 #045 #046 #047 #048 #050 #051 #052 #053 #055 #056 #057 #058 #059 #060 #061 #062 #063 #064 #065 #066 #067 #068 #069 #070 #071 #072 #073 #074 #075 #076 #077 #078 #079 #<u>080</u> #081 #082 #083 #084 #085 #086 #087 #088 #089 #090 #091 #092 #093 #094 #096 #097 #098 #099 #100 #101 #102 #103 #104 #105 #106 #107 #108 #109 #110 #111 #112 #113 #114 #115 #116 #117 #118 #119 #120 #121 #122 #123 #124 #125 #126 #127 #128 #129 #130 #131 #132 #133 #134 #135 #136 #138 #139 #140 #141 #142 #143 #144 #145 #146 #147 #148 #149 #151 #152 #153 #154 #155 #156 #157 #158 #159 #161 #170 #<u>171</u> #<u>172</u> #173 #177 #200 #<u>207</u> #209 #210 #213 #215 #216 #217 #219 #228 #229 #241 #242 #230 #237 #238 #243 #244 #245 #<u>247</u> #250 #<u>251</u> #252 #264 #265 #266 #267 #268 #269 #270 #273 #275 #276 #280 #281 | 180 | 83.85% | $\leq 3.00\text{E-}14$ | |
| Other ray-finned | #008 #<u>054</u> #137 #150 #<u>160</u> #162 #163 #<u>167</u> #174 #211 #218 #227 #232 #<u>277</u> | 14 | 77.51% | $\leq 5.00\text{E-}12$ | |
| Non-specific | #175 #178 #184 #197 #198 #<u>202</u> #<u>203</u> #<u>204</u> #206 #212 #220 #221 #222 #235 #248 #<u>278</u> | 16 | 65.45% | $\leq 3.00\text{E-}9$ | the lowest mode |

**Table S5.** Results from DNA Extraction and Sequencing.

| ID | Fossil portions | Sample weight (g) | aDNAmix (ng/mL) | Sequences amount |
| --- | --- | --- | --- | --- |
| 1 | texture | 30 | 33 | 22253 |
| 2 | texture | 28 | 54 | 30932 |
| 3 | texture | 40 | 99 | 445752 |
| 4 | texture | 50 | 141 | 735110 |
| 5 | texture | 25 | 35 | 24976 |
| 6 | non-texture | 50 | 73 | 52 |
| 7 | non-texture | 50 | 82 | 583 |

